# Towards the new normal: Transcriptomic and genomic changes in the two subgenomes of a 100,000 years old allopolyploid, *Capsella bursa-pastoris*

**DOI:** 10.1101/479048

**Authors:** Dmytro Kryvokhyzha, Pascal Milesi, Tianlin Duan, Marion Orsucci, Stephen I. Wright, Sylvain Glémin, Martin Lascoux

## Abstract

Allopolyploidy has played a major role in plant evolution but its impact on genome diversity and expression patterns remains to be understood. Some studies found important genomic and transcriptomic changes in allopolyploids, whereas others detected a strong parental legacy and more subtle changes. The allotetraploid *C. bursa-pastoris* originated around 100,000 years ago and one could expect the genetic polymorphism of the two subgenomes to become more similar and their transcriptomes to start functioning together. To test this hypothesis, we sequenced the genomes and the transcriptomes (three tissues) of allotetraploid *C. bursa-pastoris* and its parental species, the outcrossing *C. grandiflora* and the self-fertilizing *C. orientalis*. Comparison of the divergence in expression between subgenomes, on the one hand, and divergence in expression between the parental species, on the other hand, indicated a strong parental legacy with a majority of genes exhibiting a conserved pattern and *cis*-regulation. However, a large proportion of the genes that were differentially expressed between the two subgenomes, were also under *trans*-regulation reflecting the establishment of a new regulatory pattern. Parental dominance varied among tissues: expression in flowers was closer to that of *C. orientalis* and expression in root and leaf to that of *C. grandiflora*. Since deleterious mutations accumulated preferentially on the *C. orientalis* subgenome, the bias in expression towards *C. orientalis* observed in flowers suggests that expression changes could be adaptive and related to the selfing syndrome, while biases in the roots and leaves towards the *C. grandiflora* subgenome may be reflective of the differential genetic load.

## Introduction

Polyploidy, and in particular allopolyploidy, whereby a novel species is created by the merger of the genomes of two species, is considered to be a common mode of speciation in plants (1) as it induces an instant reproductive isolation, the difference in chromosome number impeding reproduction with the parental species. In the case of allopolyploidy, the daughter species thus has at inception two divergent sub-genomes, one inherited from each parental species. Such an increase in genome copy number can be advantageous and could partly explain the apparent evolutionary success of allopolyploid species (2, 3). For instance, genome doubling creates genetic redundancy, thereby increasing genetic diversity and allowing the masking of deleterious mutations through compensation. Genome doubling and initial redundancy also offer new possibilities for the evolution of genes over time: one copy can degenerate, both can be conserved by dosage compensation through, for instance, compensatory drift (4) or their pattern of expression can diverge and even lead to the evolution of new functions (see (5) and references therein). Moreover, gene redundancy also potentially allows tissue-specific expression of different gene copies (6, 7). On the other hand, as pointed out by many authors (8–11), the evolutionary success of allopolyploids can also appear paradoxical since the birth of a new allopolyploid species will also be accompanied by numerous challenges. These challenges are first associated with the initial hybridization between two already divergent genomes that have now to start working together, implying, among other things, important changes of the meiotic machinery and of gene expression patterns (12).

The magnitude of gene expression changes has been reported to vary substantially across polyploid species, from minor modifications (13, 14) to so-called “transcriptomic shock” (7). The balance in expression pattern between the two subgenomes also seems to be highly variable and ranges from non-additivity, such as extreme expression dominance of one of the ancestral genomes over the other, to the additivity of their expression contributions (15, 16), and it also evolves through time. For example, in *Mimulus peregrinus* gene expression dominance was established early on but also increased over successive generations (17). However, the generality, timing, and causes of changes in expression pattern of the two parental genomes remain poorly known beyond a few case studies (16, 18) and may, to a large extent, depend on parental legacy because a part of the observed differences between the two subgenomes of the allopolyploid species may have already been present between the parental species (3).

Ultimately, changes in patterns of gene expression will follow from modifications in gene expression regulation. Differences in gene expression can be due to changes in *cis*-and *trans*-regulatory elements. *Cis*-regulatory elements alter allele-specific expression and are generally located close to the gene they regulate (e.g., promoters), whereas *trans*-regulatory elements affect both alleles and can be located anywhere in the genome (19–22). In the case of a newly formed allopolyploid species, one would expect the two copies of a gene to be under the influence of *trans*-regulatory elements inherited from both parents and its expression level to first move towards the mean expression of the two parental species. Retaining the parental pattern of expression in each subgenome would imply that only *cis*-regulation takes place, or there are forces opposing the establishment of cross *trans*-regulation. For instance, one could expect purifying selection to have a larger impact on *trans*-acting mutations than on *cis*-acting ones because the former are more pleiotropic than the latter. If so, the residual variants will mostly be *cis*-acting ((23) but see (24)). It was also shown that a gene is often under the influence of both *trans*-and *cis*-regulatory elements that act in opposite directions (22), leading to a *cis*-*trans* compensation that prevents overshooting optimal overall expression level. Such compensation between *cis*-and *trans*-regulatory elements is one of the predictions of the enhancer runaway (ER) model proposed by Fyon et al. (25). Under the ER model, and especially in outcrossing species where heterozygotes are frequent, *cis*-regulatory variants facilitate the exposure of alleles to purifying selection. If the enhancer and the gene they regulate are linked then the up-regulating variants will hitch-hike with the allele carrying the lowest number of deleterious mutations, leading to an open-ended escalation in enhancer strength (25). As selection on expression appears to be primarily stabilizing (22, 26, 27), at least at intermediate evolutionary timescales (28), a compensatory effect of expression in *trans* is predicted (25, 29). The relative importance of *cis*-and *trans*-regulation can be examined by comparing the relative expression in the parental species with the relative expression of homeologous genes in the newly formed tetraploid (19, 30, 31).

Finally, differential expression between the two genomes could result from a differential accumulation of deleterious or slightly deleterious mutations between the two subgenomes or, alternatively, be also related to biased phenotypic or adaptive changes associated to the differences between the two parental species. If the differential expression is *only* due to differential accumulations of deleterious mutations, we would expect to see the same differential expression pattern across different tissues, whereas if it is related to biased phenotypic or adaptive changes then we may expect to see differences depending on the tissue considered.

The shepherd’s purse *C. bursa-pastoris* is an allotetraploid selfing species that originated some 100-300 kya from the hybridization of the ancestors of *C. orientalis* and *C. grandiflora* (14) (Fig. 1A). The two parental species are strikingly different: *C. orientalis* (hereafter *CO*), a genetically depauperate selfer, occurs across the steppes of Central Asia and Eastern Europe (32), whereas *C. grandiflora* (hereafter *CG*), an obligate outcrosser with a particularly high genetic diversity, is primarily confined to a tiny distribution range in the mountains of Northwest Greece and Albania (32) (Fig. 1). Among *Capsella* species, only *C. bursa-pastoris* has a world-wide distribution (32), some of which might be due to extremely recent colonization events associated with human population movements (32, 33). In Eurasia, the native range of *C. bursa-pastoris* is divided into three genetic clusters - Asia, Europe, and the Middle East (hereafter ASI, EUR and ME, respectively) - with low gene flow among them and strong differentiation both at the nucleotide and gene expression levels (33, 34). Reconstruction of the colonization history suggested that *C. bursa-pastoris* spread from the Middle East towards Europe and then expanded into Eastern Asia. This colonization history resulted in a typical reduction of nucleotide diversity with the lowest diversity being found in the most recent Asian population (33).

**Fig. 1.**
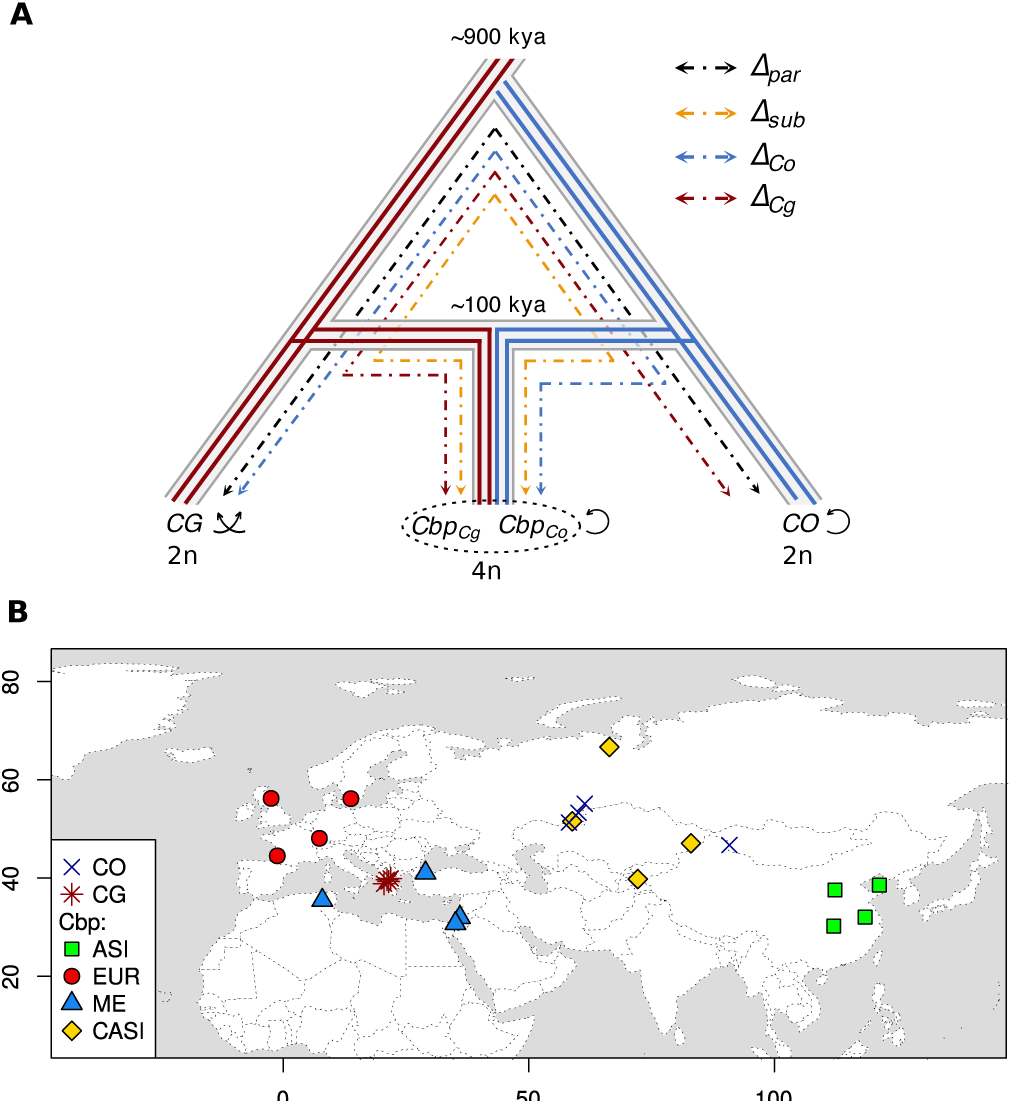
Evolutionary history and sampling locations of the three *Capsella* species used in this study. **A.** Solid lines represent subgenomes segregation after *C. grandiflora* (*CG*), and *C. orientalis* (*CO*), ancestors hybridization. Red is *C. grandiflora* genetic background and blue is *C. orientalis* genetic background. The ploidy level (n) and the reproductive system are also indicated. Dashed and dotted lines represent the comparisons used to compute the gene expression convergence index (see Material and methods). **B.** CO, CG, ASI, EUR, ME, CASI correspond to *C. orientalis*, *C. grandiflora*, and four populations of *C. bursa-pastoris*, *Cbp*, Asia, Europe, Middle East, and Central Asia, respectively. We shifted slightly population geographical coordinates when those overlapped in order to make all of them visible on the map.

It has been possible to phase the sub-genomes by assigning each genome sequence (or transcript) to a parental species sequence (35). The phased data suggested that the differences in deleterious variants between the two subgenomes of *C. bursa-pastoris* are largely a legacy of the differences between the two parental species and that biased fractionation, the biased loss of ancestral genomes in an allopolyploid, is limited (14, 36). A recent study further demonstrated that the evolutionary history of the two subgenomes varies across the different populations (35): for example, selective sweeps were more common on the subgenome descended from *C. grandiflora* (hereafter *Cbp_Cg_*) than on the subgenome descended from *C. orientalis* (hereafter *Cbp_Co_*) in Europe and the Middle East, while the opposite pattern was observed in Asia (35). There were also differences in gene expression: the two subgenomes showed no significant difference in the levels of expression in Asia, whereas the *Cbp_Cg_* subgenome was slightly more over-expressed than the *Cbp_Co_* subgenome in Europe and the Middle East. The study by Kryvokhyzha et al. (35), however, did not include expression of the two parental species and considered only the expression in one tissue (seedlings). We thus do not know yet how the current state was established since the origin of the polyploid.

The aim of the present study was to address questions on the evolution of the pattern of expression of two subgenomes of the allotetraploid shepherd’s purse *C. bursa-pastoris* since they derived from the two parental species. We focused on two main questions. First, has the relative contribution of *cis*- and *trans*-regulation been altered by polyploidization? Second, could differential expression between the two subgenomes only results from a differential accumulation of deleterious/slightly deleterious mutations between the two subgenomes (nearly neutral hypothesis) or is it *also* related to phenotypic differences between the two parents (adaptive hypothesis)? Since one parent is outcrossing (*C. grandiflora*) and the other self-fertilizing (*C. orientalis*), the former with large flowers and the latter with tiny ones, one may expect differential expression in flower tissues of selfing *C. bursa-pastoris* to be biased towards the *C. orientalis* expression levels under the adaptive hypothesis whereas tissues that have not experienced adaptive specialization might show an expression bias towards *C. grandiflora*.

To address these questions and, more generally, to characterize the expression pattern of *C. bursa-pastoris*, we analyzed the genomes and the transcriptomes of three tissues (flowers, leaves, and roots) of 16 accessions coming from different populations of the *C. bursa-pastoris* natural range and compared them with those of the parental lineages *C. grandiflora* and *C. orientalis* (four accessions each). In total, 72 transcriptomes and 24 genomes were analyzed.

One hundred thousand generations after its inception, *C. bursa-pastoris* does not show any sign of a transcriptomic shock. Instead, our data revealed highly concerted changes with the expression levels of the two subgenomes converging towards a median value. This was achieved by a balance between *cis*-and *trans*-regulation and a strong parental legacy that was also observed for the accumulation of deleterious mutations over the two subgenomes. While the differential accumulation of deleterious mutations between subgenomes could explain part of the differential expression between them there were also significant tissue-specific differences in subgenome dominance and convergence, indicating that adaptive changes may also have contributed to the evolution of the expression patterns of the two subgenomes.

## Material and methods

### Samples, sequencing and data preparation

We obtained the whole genome and RNA-Seq data from flower, leaf and root tissues of (i) 16 accessions of *C. bursa-pastoris* coming from already characterized populations from Europe (EU), the Middle East (ME) and Eastern Asia (ASI) (33) and from hitherto unstudied Central Asian populations (CASI) and (ii) four accessions each of *C. grandiflora* and *C. orientalis* (Fig. 1). The genomic data included both published and newly sequenced genomes (Table S1). For newly sequenced genomes, DNA was extracted from leaves with the Qiagen DNeasy Plant Mini Kit, libraries were prepared using the TruSeq Nano DNA kit, and 150-bp paired-end reads were sequenced on Illumina HiSeqX platform (SciLife, Stockholm, Sweden). All 72 RNA-Seq libraries (24 accessions×three tissues) were sequenced in this study. For RNA sequencing, seeds were surface-sterilized and germinated as dexsscribed in (34). Seedlings were then transplanted into pots (10×10×10cm) filled with soil seven days after germination and cultivated in one growth chamber (22°C, 16:8h light/dark period, light intensity 150 *µmol/m*^2^*/s*). Seven days after the onset of flowering, flower buds, leaves, and roots were collected, snap-frozen in liquid nitrogen, and stored at −80°C before extraction following manufacturer protocol (Plant Total RNA Kit (Spectrum) for flower buds and leaves, and RNeasy Plant Mini Kit (Qiagen) for roots). RNA sequencing libraries were prepared using the TruSeq stranded mRNA library preparation kit including polyA selection and sequenced for 125-bp paired-end reads on Illumina HiSeq 2500 platform (SciLife, Stockholm, Sweden). Sequencing of new samples yielded an average library size of 57 million reads for DNA sequencing and 59 million reads for RNA-Seq.

DNA and RNA-Seq reads were mapped to the *C. rubella* reference genome (37) with Stampy v1.0.22 (38). To account for the divergence from the reference genome, the substitution rate was set to 0.025 for *C. bursa-pastoris*, 0.02 for *C. grandiflora*, and 0.04 for *C. orientalis*. On average, 85%, 90% and 85% of the DNA reads were successfully mapped for the corresponding three species and 98% in all species for RNA mapping. This yielded an average coverage of 51x and 52x for DNA and RNA data, respectively. Genotyping of DNA and RNA-Seq alignments was performed using HaplotypeCaller from the Genome Analysis Tool Kit (GATK) v3.5 (39) as described in (35). The subgenomes of *C. bursa-pastoris* were phased with HapCUT version 0.7 (40) following the procedure by (35). The quality of this phasing procedure was ascertained by comparing the phased subgenomes with the subgenome assembly obtained by (36). The un-phased expression data was generated for non-overlapping feature positions (option: *-m union*) using the *htseq-count* program from HTSeq v0.6.1 (41). To compare the expression between the two subgenomes of *C. bursa-pastoris*, homeologue-specific counting of alleles was performed using *ASEReadCounter* from GATK and phased according to the phased genomic data. We analyzed only the counts of SNPs that showed no strong deviation from the 0.5 mapping ratio in DNA data defined with a statistical model developed by (42). To make the homeologue-specific count data of *C. bursapastoris* comparable with parental read count data (allelic counting underestimates the expression of genes with a low number of heterozygous sites), we scaled the homeologue-specific counts using the unphased data and the allelic ratio in the phased data.

### Population structure

In order to assess the relationship of the newly obtained Central Asian samples with other populations, we characterized population structure through a principal component analysis (*ade4* R package (43)) and by reconstructing a phylogenetic tree from the genomic Single Nucleotide Polymorphisms (SNPs) of the 24 accessions (neighbor-joining algorithm on absolute genetic distance, *ape* R package (44)). Relationships between samples were also explored in the expression data using a principal component analysis (*ade4* R package (43)) and hierarchical distance clustering with bootstrap support (*pvclust* R package, (45)).

### Gene expression analyses

Given that the gene expression patterns in homeologue-specific and total expression can produce different results (46), we performed the differential gene expression analyses on both the unphased data and phased data. We first assessed the differences between populations of *C. bursa-pastoris* and parental species in unphased data by partitioning the analysis between tissues and populations. In a second step, the phased data was used to assess homeologue-specific expression differences between subgenomes in different tissues and populations. Finally, we analyzed the differences in expression patterns in parental diploid species and *C. bursa-pastoris* by classifying expression patterns into categories in both phased and unphased data.

Differential gene expression analyses were carried out in *edgeR* (47). The TMM normalization for different library sizes (47) was used for differential gene expression analyses, while for all other analyses, we used the count per million (CPM) normalization (one was added to every gene count to bypass log-transformation of zero expression). Phased counts were normalized by the mean library size of the two subgenomes 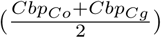 and only genes showing no strong mapping bias were retained (see below). For both datasets (unphased or phased), only genes with at least one sample having a non-zero expression in every population/species were kept.

Differences between the two subgenomes (homeologue-specific expression were assessed with the integration of the information from both RNA and DNA data to exclude highly biased SNPs and to account for the noise in read counts due to statistical variability. The data were analyzed using the three-stage hierarchical Bayesian model for allelic read counts developed by (42). The model was implemented using Markov chain Monte Carlo (MCMC) with 200,000 iterations with burn-in of 20,000 and thinning interval of 100. Each analysis was run three times to assess convergence. The significance specific expression (HSE) was defined from a Bayesian analog of the false discovery rate (*FDR* < 0.05). Expression patterns in *C. bursa-pastoris* and its parental species were classified into categories based on significant and non-significant differential expression defined with *edgeR* (47). We considered the four genomes/subgenomes (*CG*, *CO*, *Cbp_cg_*, and *Cbp_co_*) and three possibilities for each of the six pairwise comparisons (significantly over, under or equally expressed, *FDR* < 0.05), and grouped the resulting combinations into seven main categories: *No difference*, *Intermediate*, *Legacy*, *Reverse*, *Dominance*, *Compensatory drift*, and *Transgressive* (see the results for categories description). We also performed similar analysis for the un-phased total *C. bursa-pastoris* expression (thus considering only three pairwise comparisons) by classifying the expression patterns into four major categories: 1) *no differential expression*, when no significant differences are detected in any of the three pairwise comparisons, 2) *intermediate*, when the expression of *C. bursa-pastoris* (*Cbp*) is intermediate between *C. grandiflora* (*CG*) and *C. orientalis* (*CO*), 3) *dominance* of one of the parents over the other, when the mean expression of *C. bursa-pastoris* is equal to only one parental species and the two parents are significantly different, and finally 4) *transgressive*, when the mean expression of *C. bursapastoris* is outside the range of expression of the two parents and significantly different from at least one parent.

### Similarity and Convergence indices

To quantify the similarity between each subgenome expression level and the expression level in the parental species, we developed a similarity index (*S*). For each transcript *i* and each subgenome *j*, *S* was computed as the subgenome relative expression deviation from the mean expression level (*E*) in the parental species, *µ_i_* = (*E_iCO_* + *E_iCG_*)/2:

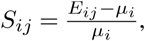

This index is centered on 0, so that if *S_ij_* < 0 or *S_ij_ >* 0, the expression of a given transcript in a given subgenome is more similar to the expression of that transcript in *CG* or *CO*, respectively. The difference between the absolute values of median *S_i_* for *Cbp_Cg_* and *Cbp_Co_* was used as a measure of dominance between the subgenomes, ∆_*S*_= |*S_CbpCo_*| − |*S_CbpCg_*|.

Finally, for each gene that was differentially expressed between the two parental species, a convergence index, *C*, was computed from the absolute difference in expression for:

- subgenomes: 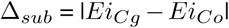
- parental species: 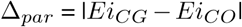
- each subgenome and the *opposite* parental species: 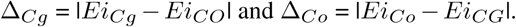

These differences correspond to the phylogenetic distances (Fig. 1A). In principle, if the regulation of gene expression in *Cbp_Cg_* is independent of the regulation of gene expression in *Cbp_Co_*, then the overall ∆_*sub*_, ∆_*par*_, ∆_*Cg*_and ∆_*Co*_are expected to be equal. To compare these quantities, for each transcript *i*, we used a convergence index (*C_i_*):

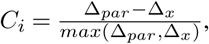

So, *C_Cbp_* measures the expression convergence of *Cbp_Cg_* toward *Cbp_Co_*, *C_CbpCo_*measures the expression convergence of *Cbp_Co_* toward *Cbp_Cg_*, and *C_Cbp_* measures the overall subgenomes convergence within *Cbp*. ∆_*x*_stands for either ∆_*Co*_, ∆_*Cg*_or ∆_*sub*_, respectively. *C_i_* thus ranges from −1 to 1, with positive values indicating more similar expression between the subgenomes of *C. bursa-pastoris* than between parental species, and negative values indicating increased differences between subgenomes; the closer *C_i_* to 0, the more similar are the expression patterns to parental species.

### Gene ontology enrichment test

For various lists of genes of interest detected in the analyses described above, gene ontology (GO) enrichment tests were performed using the *topGO* R package (48). The GO term annotation was downloaded from PlantRegMap (http://plantregmap.cbi.pku.edu.cn/download.php#go-annotation) and used as a reference set for *topGO* (i.e., custom input). Fisher’s exact-test procedure (*weight* algorithm) was performed to assess the enrichment (*p* < 0.05) for either molecular functions (MF) or biological processes (BP). Finally, the *REViGO* software (49) was used to remove GO terms redundancy and to cluster remaining terms in a two-dimensional space derived by applying multidimensional scaling to a matrix of the GO terms semantic similarities. Cytoscape v3.6.1 was used to visualize GO terms networks (50).

### Difference between species and subgenomes in dele-terious mutations

To compare the number of deleterious mutations between the two subgenomes of *C. bursa-pastoris*, we classified mutations into tolerated and deleterious ones (DEL) using SIFT4G (51). We used *C. rubella* (35) and *Arabidopsis thaliana* (TAIR10.22) SIFT4G reference databases. This helps avoid reference bias towards *C.rubella* away from calling mutations to be deleterious in the *C. grandiflora* homeologue. We considered only the mutations that accumulated after speciation of *C. bursa-pastoris* and identified mutations specific to *C. grandiflora*, *C. orientalis*, the two subgenomes of *C. bursa-pastoris*, and *Neslia paniculata* that was used as an outgroup here. All estimates were relative to the total number of SIFT4G annotated sites to minimize the bias associated with variation in missing data as in (35). Only the European and Middle Eastern populations were used in further analysis of the distribution of deleterious mutations, in order to exclude the effect of gene flow between *C. orientalis* and the Asian population of *C. bursa-pastoris* (35).

We assessed the distribution of deleterious mutations between the two subgenomes of *C. bursa-pastoris* to test whether they accumulated (i) more in one gene copy than in the other at the homeologue level, as would be expected under a pseudogenization process, (ii) more in one subgenome than in the other as expected if one subgenome predominates. Under the null hypothesis (random accumulation without subgenome bias) the distribution of deleterious mutations between the two subgenomes should follow a binomial distribution with mean 1/2. Under the first hypothesis, the distribution should be more dispersed with the same mean, which can be modeled by a Beta-binomial distribution. Under the second hypothesis, the mean should differ from 1/2. However, over-dispersion and bias can also occur because of missing data and sampling error, we thus used synonymous mutations (SYN) to control for this and built the correct null distribution. To do so, we developed a maximum likelihood method implemented in *R* (52) as follows. First, we identified a most likely probability distribution model by fitting four models to the SYN dataset, where *nSY N* is the sum of *SY N* mutations occurring on both homeologous genes and *kSY N* is the number of *SY N* mutations occurring on *Cbp_Cg_* genes. The four models are:

- M1: *kSY N ~ B*(*nSY N*, 0.5), a binomial distribution with no bias between *Cbp_Cg_* and *Cbp_Co_*,
- M2: *kSY N ~ B*(*nSY N*, 0.5 + *b*), a binomial distribution with bias,
- M3: *kSY N ~ BB*(*nSY N*, 0*, φ*), a beta-binomial distribution with no bias,
- M4: *kSY N ~ BB*(*nSY N, b, φ*), a beta-binomial distribution with bias.

For convenience, the beta-binomial distribution:

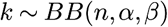

was re-parameterized as:

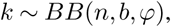

where 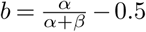 and 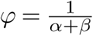 (53, 54). In this way, the parameter *b* was a measure of the bias towards the *Cbp_Cg_* genes, and *φ* was a measure of the variance of the probability that a mutation is found within the *Cbp_Cg_* homeologues, and can be interpreted as an index of overdispersion. A large value of *φ* indicates that mutations tend to accumulate preferentially in one of the two homeologous genes, and a small value of *φ* indicates that mutations are more evenly distributed between them. We calculated the likelihood of each model and chose the best-fitting model with a hierarchical likelihood ratio test (hLRTs).

After choosing the beta-binomial distribution with bias as the most likely null distribution, we estimated the parameters *b* and *φ*. We introduced a new set of models to test for the specific features of the distribution of deleterious mutations:

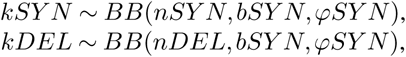

The null model assumes that both parameters *b* and *φ* are the same for the *SY N* and *DEL* datasets, while the alternative models allow the *DEL* dataset to have different parameters from the *SY N* dataset: only *bDEL*, only *φDEL*, or both *bDEL* and *φDEL* were allowed to vary. We calculated the likelihood of each model, chose the best fitting model with hierarchical likelihood ratio tests (hLRTs) and estimated the parameters of the selected model. Bootstrap estimates of confidence intervals were estimated with 1000 bootstrap replicates.

### Relationship between deleterious mutations and gene expression

The SIFT4G annotation of the *C. rubella* database was used to match the gene IDs of the mutation and expression data. For each tissue, the relationship between the bias in the number of deleterious mutations between subgenomes and the bias in homeologue expression was investigated by calculating, for gene *i* in accession *j*, the difference (*d_ij_*) in the number of deleterious mutations (*DEL*) between homeologous gene pairs:

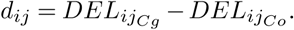

The expression ratio between the homeologues of genes with significant HSE was used as a measure of homeologue expression bias:

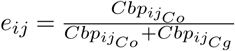

Genes were further classified into four categories according to the deleterious mutations bias, *d*, and homeologue expression bias, *e*:

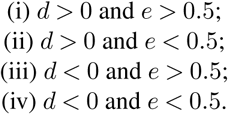

Genes with no bias in the distribution of deleterious mutation (*d* = 0) or no significant HSE (*FDR* < 0.05) were removed from the analysis. Fisher’s exact test was then used to test for independence between the difference in the number of deleterious mutations (*d*) and homeologue expression bias (*e*). As a control, the whole analysis was reproduced with *d_ij_* computed from the number of synonymous mutations in genes with no *DEL* mutations. In addition, we also compared the number of silenced genes (genes with zero expression values) of each subgenome of *C. bursa-pastoris*, to check if there was a relationship between genetic load and silenced genes.

## Results

### Population genetic structure

The SNP-based PCA (670K genomic SNPs without any missing data) confirmed the phylogenetic relationships between *C. grandiflora* (*CG*), *C. orientalis* (*CO*) and *C. bursa-pastoris* (*Cbp*) described in (33– 35). The first principal component (PC) explained most of the variance (66%) and clearly discriminated *CG* and the *Cbp_Cg_* subgenome from *CO* and the *Cbp_Co_* subgenome (Fig. 2, top-left panel). To investigate further the population structure within *C. bursa-pastoris*, we then focused on genetic variation in each subgenome (Fig. 2, middle and bottom-left panels, respectively for *Cbp_Cg_* and *Cbp_Co_*). In both cases, there were three main clusters gathering accessions from Europe (EUR), Asia (ASI), and the Middle East (ME), respectively. Accessions from Central Asia (CASI) tended to cluster with European accessions for both subgenomes, even if they were more scattered (especially for the *Cbp_Cg_* subgenome). A phylogenetic analysis also confirmed that the new samples from Central Asia were most similar to the European genetic cluster and showed that they did not form a separate genetic cluster (Fig. S1).

**Fig. 2.**
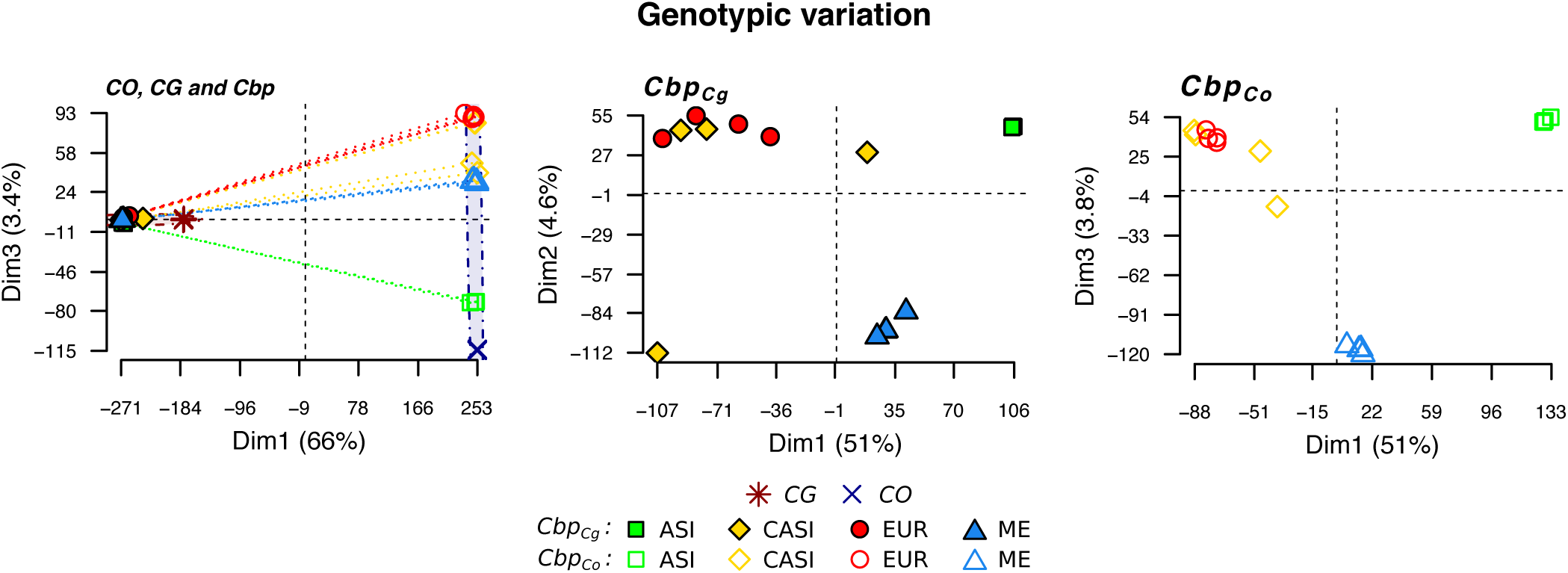
Genomic variation patterns in three *Capsella* species. Variation was visualized with a principal component analyses based on the SNPs of *C. grandiflora* (*CG*), *C. orientalis* (*CO*), and *C. bursa-pastoris* (*Cbp*): Asian (ASI), Central Asian (CASI), European (EUR), and Middle Eastern (ME) populations for the three species (top panel) or only for the subgenomes of *C. bursa-pastoris* (*Cbp_Cg_* and *Cbp_Co_*, middle and bottom panels).

### Global variation in gene expression reflects genetic relationships

Pairwise comparisons of a number of differentially expressed (DE) genes between species (unphased data, 16,039 genes) showed that patterns of expression varied across tissues. First, the number of differentially expressed genes between the parental species was the highest in flower tissues, while leaf tissues were the least differentiated (Table S2). Second, in flowers, overall gene expression of *C. bursa-pastoris* was the closest to *C. orientalis*, while in the two other tissues it was the closest to *C. grandiflora* (Table S2). At the population level, no clear pattern appeared: for instance, ME accessions were the closest to *C. grandiflora* in roots, while ASI accessions were the closest to *C. grandiflora* in leaves and CASI accessions in flowers (Table S3).

Gene expression variation was then surveyed in 11,931 genes for which phased expression of the two subgenomes was available in all populations of *C. bursa-pastoris*. Clustering of population/species mean expression values confirmed that the main difference in overall expression variation was between tissues (Fig. S3). The principal component analyses of the three tissues separately (Fig. 3) revealed that the global variation pattern in gene expression reflected phylogenetic relationships (Fig. 3 and Fig. S2). The two subgenomes of *C. bursa-pastoris* were most similar to their corresponding parental genome along the Dim1, *i.e.* expression in the *Cbp_Cg_* subgenome grouped with *C. grandiflora*, and the *Cbp_Co_* subgenome grouped with *C. orientalis*. The Dim2 reflected population structure; here again CASI accessions grouped with EUR accessions.

**Fig. 3.**
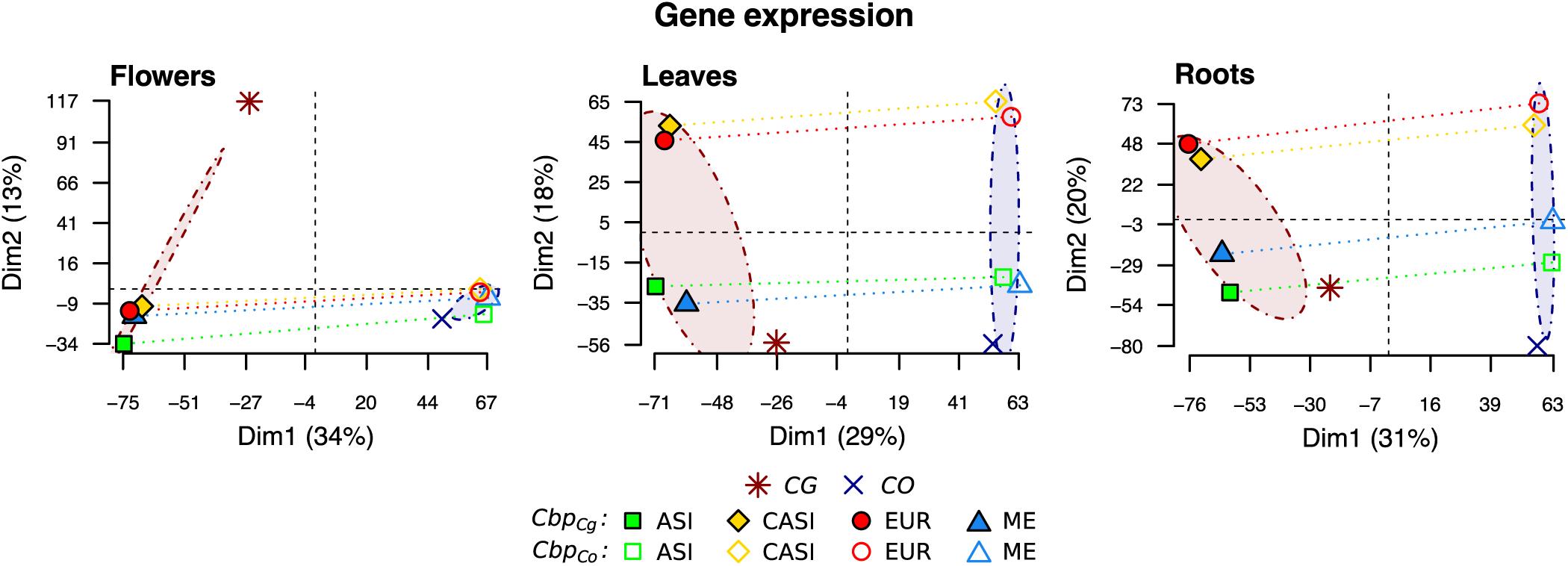
Transcriptomic variation patterns in three *Capsella* species. Variation was visualized with a principal component analyses of phased gene expression data (11,931 genes) for the three different tissues. CO, CG, ASI, EUR, ME, CASI correspond to *C. orientalis*, *C. grandiflora*, and four populations of *C. bursa-pastoris*, *Cbp*, Asia, Europe, Middle East and Central Asia, respectively.

Testing for homeologue-specific expression (HSE) in *C. bursa-pastoris* showed that on average 4,096 genes (~34%) per sample were significantly differentially expressed between the two subgenomes (*FDR* < 0.05). The expression ratio between subgenomes (defined as 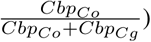 was on average 0.496 across all genes and 0.493 across genes with significant HSE indicating no strong bias towards one of the subgenomes (Table S4). The ratio in DNA reads was 0.497 and thus there was no strong mapping bias towards either subgenome. Analyses of differential expression revealed no bias in the number of differentially expressed genes toward one subgenome either when comparing tissues (Table S5A, flowers and leaves being the most differentiated tissues and leaves and roots the least) or *Cbp* populations (Table S5B, Middle East and Asia being the most distant, except for *Cbp_Co_* in flowers, while Europe and Central Asia are the closest).

### Strong parental legacy and both *cis*- and *trans*-regulatory changes

In order to investigate the total expression level changes in *C. bursa-pastoris* after *C. grandiflora* and *C. orientalis* hybridization, expression patterns of unphased data across the three species were classified into four categories: *No difference*, *Intermediate/Additivity*, *Dominance* and *Transgressive* (Table 1). Up to 55-80% of the genes in *C. bursa-pastoris* were expressed at the same total level as in the parental species and 5 to 10% showed levels of expression intermediate to that of parental species. The dominance of one parental species over the other was most evident in flowers and roots. In flowers, ~14% of *C. bursa-pastoris* genes were expressed at the same level as in *C. orientalis* but differed significantly from *C. grandiflora*, and ~8% were expressed at the same level as in *C. grandiflora* but at a different level than in *C. orientalis*. The opposite dominance pattern was detected in the root tissue. Finally, a transgressive expression pattern, when expression levels in *C. bursa-pastoris* exceeded or were lower than the expression level of both parents, was detected in 8-16% of genes.

**Table 1.**
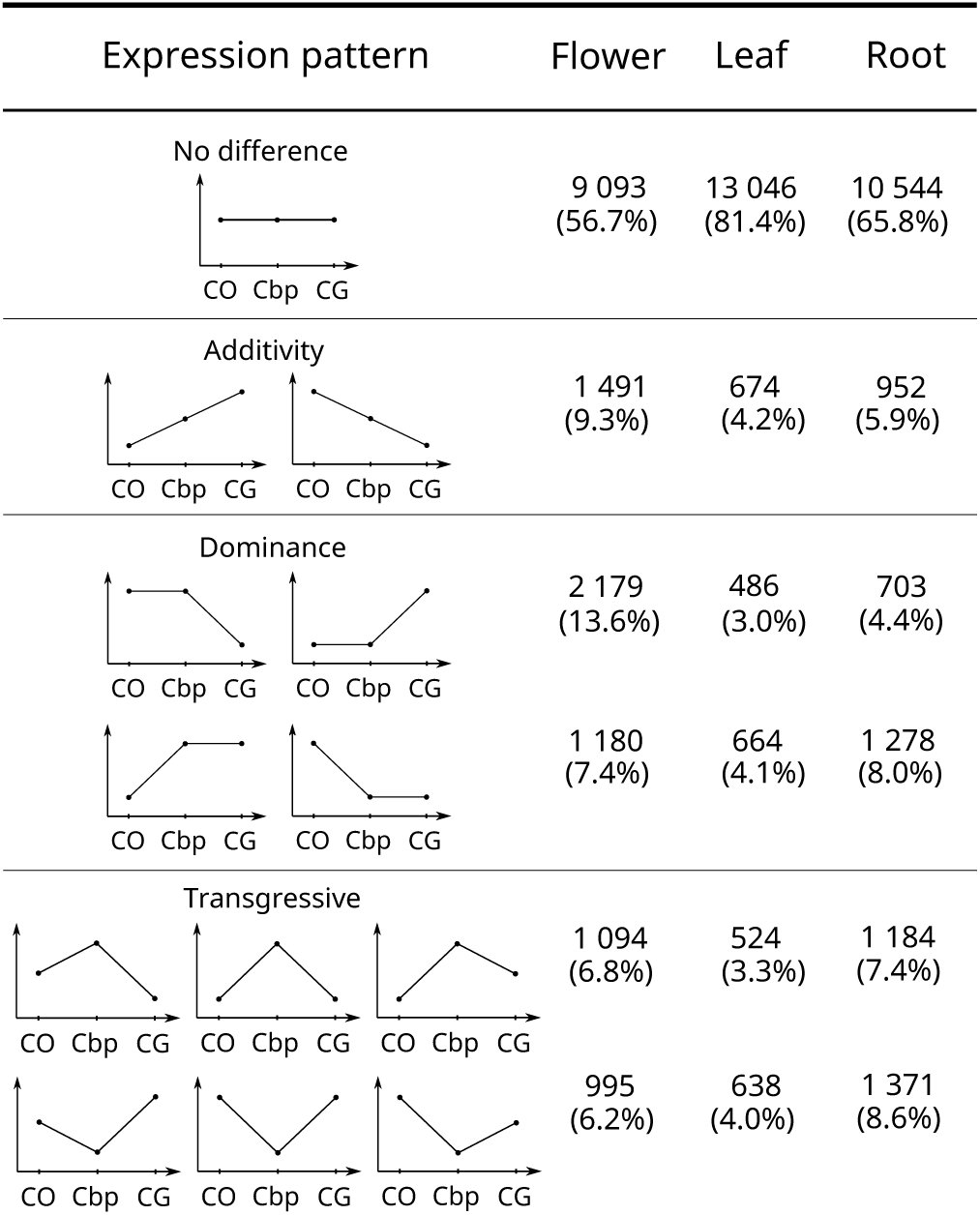
Levels of gene expression in *C. bursa-pastoris* relative to its parental species. CO, CG, and Cbp correspond respectively to *C. orientalis*, *C. grandiflora*, and *C. bursa-pastoris*. The y-axis is the level of expression, they were considered as significantly different for *FDR* < 0.05. In total, 16,032 genes were analyzed.

Gene expression in *C. bursa-pastoris* was further investigated by assessing the relative importance of *cis-* and *trans*-regulatory elements. The expression ratio of the two subgenomes was compared to the expression ratio between the two parental species (Fig. 4A). For a given gene, if its expression in the homeologous genes of *C. bursa-pastoris* is only regulated by *cis*-regulatory changes, it should be completely explained by the divergence between the parental species (the diagonal line in Fig. 4A). On the other hand, if homeologous genes are equally expressed in *C. bursapastoris* but not in the parental species, this means that *Cbp* expression is mainly controlled by *trans*-regulatory elements (the horizontal line in Fig. 4A) (19). First, the relationship between expression ratios in *C. bursa-pastoris* and parental species was positive and highly significant for all three tissues (*p* < 0.001), and the slope was intermediate between what would be expected if there were either only *cis*- or only *trans*-regulatory changes (*β* = 0.37, 0.42 and 0.46, respectively for flowers, leaves and roots). This indicates a strong parental legacy effect in expression of the two subgenomes of *C. bursa-pastoris* and suggests a joint effect of *cis*- and *trans*-regulation. Second, the variance of the expression ratio between subgenomes was significantly smaller than the variance of the expression ratio between parental genomes (Fisher’s variance test, all *p* < 0.001), indicating that the two subgenomes are closer to each other than the parental genomes are, therefore supporting a co-regulation of the two subgenomes through a mixture of *trans*- and *cis*-regulation (19, 30). Finally, the slope of the regression between the two expression ratios was the weakest in flowers, suggesting a slightly stronger *trans-*regulation and a higher level of constraints in this tissue than in roots and leaves (30).

**Fig. 4.**
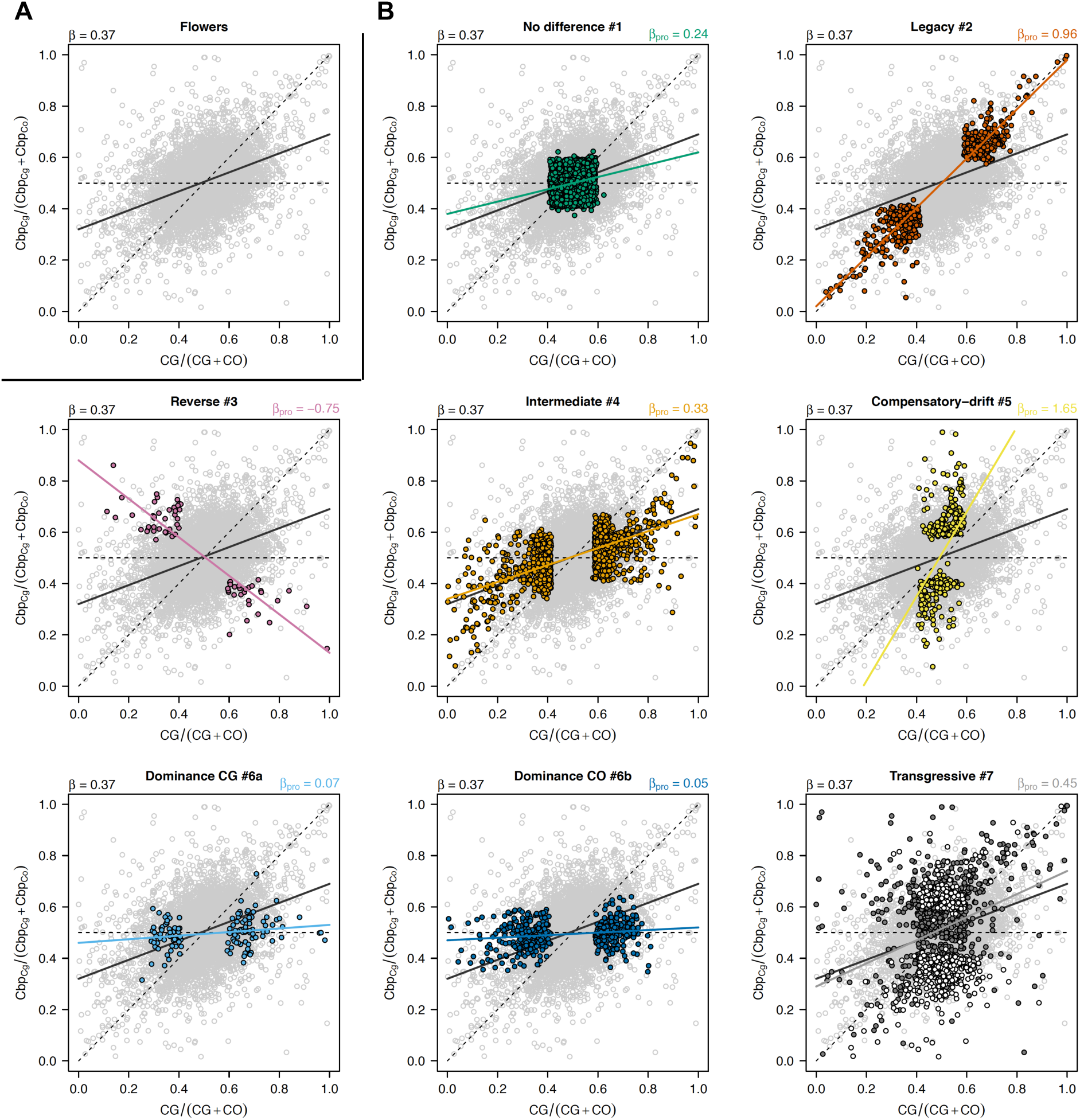
Relationships between the relative expression between the *C. bursa-pastoris* subgenomes and the relative expression between parental species. The figure shows expression in flower as an example.**A** Top-left panel is for all transcripts (11,931), **B** transcripts belonging to a specific category are colored in the others panels. The diagonal dashed lines indicate 100% *cis*-regulation divergence while the horizontal dashed lines indicate 100% *trans*-regulation. The solid lines give the slopes of the linear regressions between both ratios either for all transcript (black) or for transcript belonging to a specific category. *β* is the slope of the corresponding regression. For *Transgressive* category (bottom right panel), dark gray corresponds to categories #7a and b, light grey is for category #7c (see Fig. 5).

As mentioned above, subgenome expression level relative to parental species expression can help to disentangle the role of *cis*- and *trans*-component on overall gene regulation. We thus classified the expression patterns between the two subgenomes and parental species in seven main categories (see Fig. 5, an example for flower tissues). The majority of the transcripts was not differentially expressed between parental genomes and subgenomes (*No difference* category), ranging from 60% in flowers to 81% in leaves (Table 2).

**Table 2.** Expression variation of *C. bursa-pastoris* subgenomes relative to expression in parental species across different tissues. The percentage of transcripts within each category is given for all genes or only differentially expressed genes (*i.e*, without *No difference* category) as well as the slope of the regression of relative expression between subgenomes and relative expression between parental species for all genes per category (*β*, see Fig. 4). The percentage of transcript showing a dominance of either *Cbp_Cg_* or *Cbp_Co_* are given in parenthesis.

| Categories |  | Flowers |  |  | Leaves |  |  | Roots |  |  |
| --- | --- | --- | --- | --- | --- | --- | --- | --- | --- | --- |
| | | Transcript (%) | DE only | $\beta$ | Transcript (%) | DE only | $\beta$ | Transcript (%) | DE only | $\beta$ |
| No diff. | 1 | 60.4 | - | 0.24 | 78.4 | - | 0.32 | 67.6 | - | 0.32 |
| Legacy | 2 | 4.5 | 11.4 | 0.96 | 2.3 | 10.6 | 0.94 | 3.2 | 9.9 | 0.96 |
| Reverse | 3 | 0.5 | 1.3 | -0.75 | 0.2 | 0.9 | -0.78 | 0.4 | 1.2 | -0.78 |
| Intermediate | 4 | 12.6 | 31.9 | 0.33 | 6.8 | 31.5 | 0.44 | 8.8 | 27.3 | 0.41 |
| Comp. drift | 5 | 3.8 | 9.7 | 1.65 | 1.5 | 6.9 | 1.52 | 2.8 | 8.7 | 1.81 |
| Dominance | 6a | 1.2 | 3.0 (24) | 0.07 | 1.7 | 7.9 (54) | 0.18 | 2.7 | 8.4 (66) | 0.14 |
|  | 6b | 3.8 | 9.6 (76) | 0.05 | 1.4 | 6.5 (45) | 0.11 | 1.4 | 4.3 (34) | 0.1 |
| Transgressive | 7a | 5.1 | 12.9 | - | 2.1 | 9.7 | - | 4.2 | 13 | - |
|  | 7b | 5.2 | 13.2 | - | 2.6 | 12 | - | 4.4 | 13.7 | - |
|  | 7c | 2.8 | 7.1 | 0.45 | 3 | 13.9 | 0.55 | 4.3 | 13.4 | 0.61 |
| Total |  | 100 | 100 | 0.37 | 100 | 100 | 0.42 | 100 | 100 | 0.46 |

**Fig. 5.**
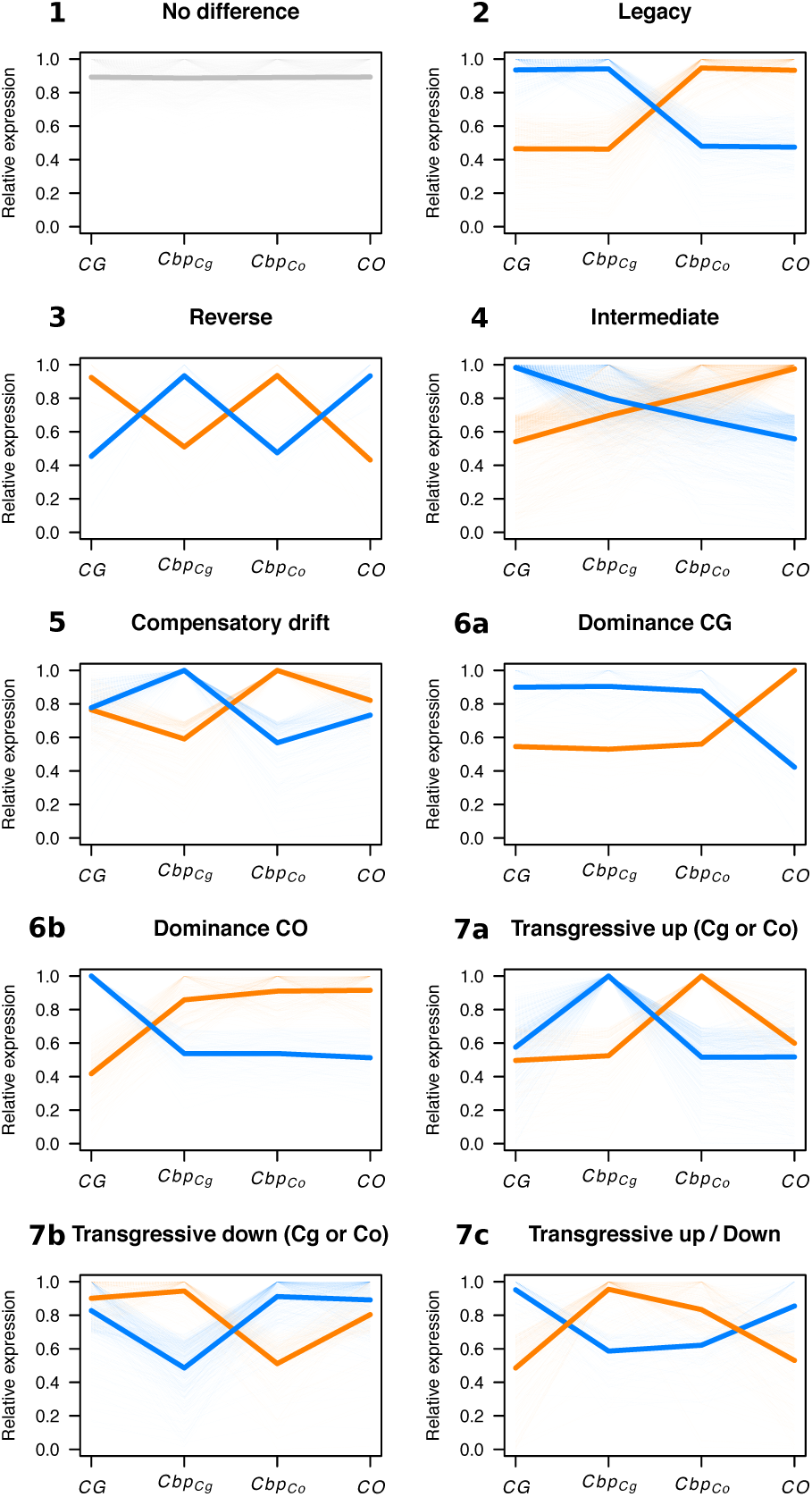
Main categories of expression variation of *C. bursa-pastoris* subgenomes relative to expression in parental species. The figure shows expression in flower as an example. Each transcript was assigned to one of seven main categories defined from its relative expression pattern across *Cbp* subgenomes (*Cbp_Cg_* and *Cbp_Co_*) and parental species (*CG* and *CO*). For each category, dashed lines correspond to single transcript relative expression to the maximal expression of this transcript in parental genomes or subgenomes and solid lines are the average expression for each genome or subgenome. Colors discriminate alternative patterns in the same category.

However, the slope of the regression between relative expression of subgenomes and parental species clearly indicated that, even if the expression levels were not significantly different between parental species and *C. bursa-pastoris* subgenomes, crossed *trans*-regulation tended to make the two subgenomes expression closer to each other than to either parental species (Fig. 4B “*No difference*” and Table 2). About 9% of genes had an *Intermediate/Additive* expression, *i.e.*, the expression of both sub-genomes being in between the expression of the two parental species. As expected this pattern was due to a combination of both *cis-* and *trans*-regulation (*β* 0.3 0.4). Only 3% showed a strict *legacy* of parental species expression which is primarily due to *cis*-regulation (*β* 1). About 4% of the genes showed a *Dominance* pattern of either *CG* or *CO* genetic background (categories 6a and 6b, Fig. 5) but the relative proportion of each background varied largely among tissues: in flowers, 76% of the transcripts showed a dominance of *CO*, while there were only 45% and 34% genes with the dominance of *CO* in leaf and root tissues (Table 2). This pattern seems to be due to a dominance of transcription factors from one subgenome over the other (*β ≈* 0.5 0.1); in favor of *CO* genetic background in flowers and *CG* in leaves and roots (Fig. 4B and Table 2). Finally, 3% of the genes had a *Compensatory-drift* profile (parental species expressions are similar but subgenome expressions diverge), a mere 0.4% showed a *Reverse* profile (each subgenome expression is similar to the opposite parental species) and about 10% of the transcripts showed a *Transgressive* pattern, either because of one (categories 7a and 7b) or of both subgenomes expression (category 7c) (Fig. 4B and Table 2). These last profiles are less straightforward to interpret in terms of *cis*-and *trans*-regulation pattern as they involve more complex post-hybridization regulation processes.

Finally, if the relative proportion of the different categories were globally conserved across tissues (Table 2), expression patterns of individual genes were strongly tissue-specific. In our data, only half of the genes showed the same expression pattern in all the three tissues. The most conserved category was *No difference* 77% and the least conserved one was *Compensatory-drift* 3%. Pairwise comparisons between tissues revealed that the number of genes with expression pattern changed between tissues was the largest between flowers and roots tissues (42%) and the smallest between leaf and root tissues (33%).

To conclude, only about 10% of the 11,931 transcripts had a transgressive or a reverse expression pattern. Expression patterns were poorly conserved between tissues except for the *No difference* category, indicating that the evolution of expression regulation is highly tissue-specific. Flower tissue differed the most from the two other tissues. In addition to a lower proportion of differentially expressed genes, flower tissues also had the lowest proportion of *Transgressive* category in the differentially expressed genes, indicating that when expression changes occurred, they either took place within the expression range of the parental species or they were compensated by the other subgenome (*Compensatory-drift*). This suggests a higher level of constraints on gene expression in flower tissues than in leaves and roots. Moreover, in flowers, the *CO* genetic background clearly dominates over the *CG* background, in striking contrast with the dominance of the *CG* genetic background in the other two tissues. Finally, expression profiles are more conserved between leaves and roots than between flowers and roots.

### Expression similarity and convergence between subgenomes: flowers differ from roots and leaves

To understand better the joint dynamics of expression in the two subgenomes across tissues, we defined a new similarity index, *S*, that measures the relative expression deviation of a given subgenome from the average parental expression. *S* indices of both subgenomes were systematically biased towards the corresponding parental genome, *i.e. Cbp_Cg_* towards *CG* and *Cbp_Co_* towards *CO* (binomial test, all *p* < 0.001) but the strength of this bias differed between subgenomes and across tissues (Fig. 6A). The distributions of *S* values for leaf and root tissues were more spread than the distribution for flowers, meaning that the relative expression in the two sub-genomes was globally less constrained in these tissues than in the flower tissue (Fig. S4).

**Fig. 6.**
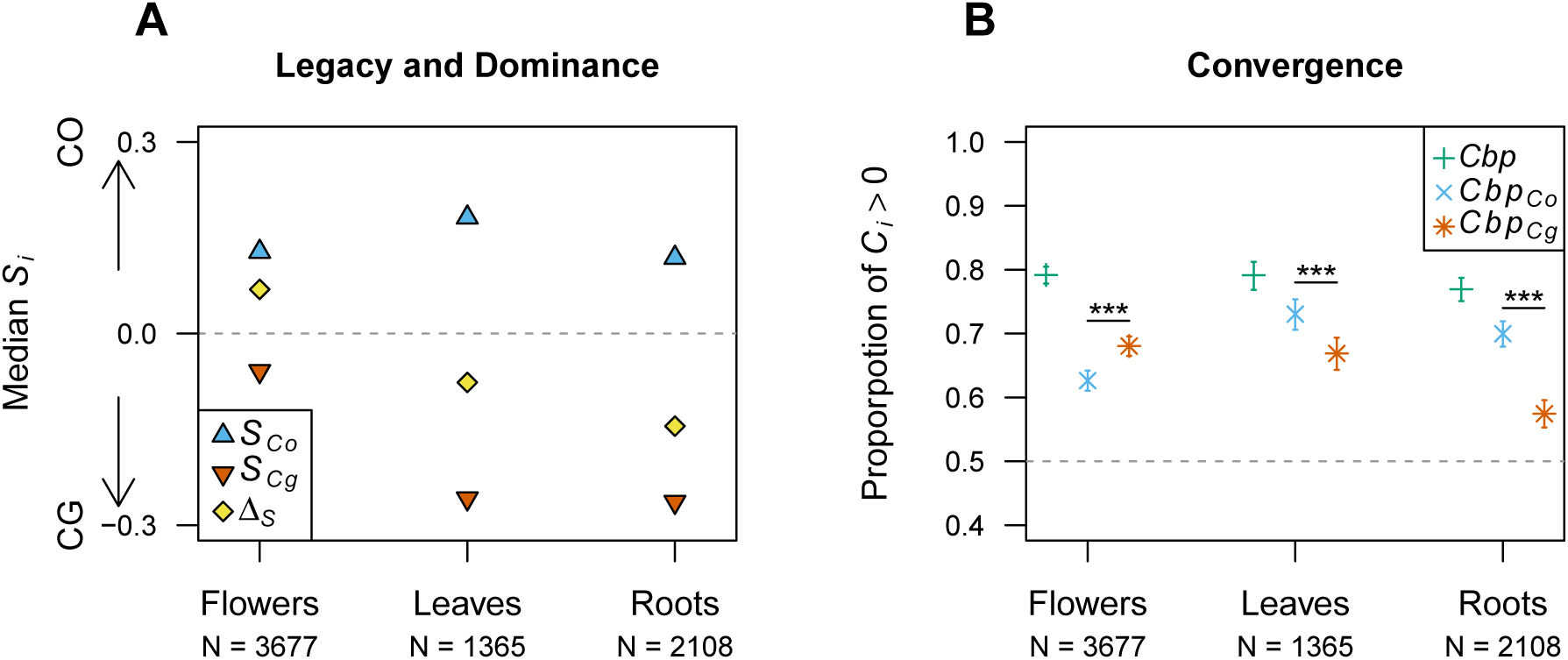
Similarity and convergence indices for differentially expressed genes between subgenomes of *C. bursa-pastoris*. **A.** For each tissue and each subgenome, the median of similarity indices for each subgenome (*S_Co_* and *S_Cg_*) are presented as well as the difference between the two indices (∆_*S*_) that indicates subgenomes dominance. Grey dotted line (*S* = 0) means no bias. **B.** The proportion of transcripts showing convergence (*C_i_>* 0) is reported for the whole genome (green plus signs) or each subgenome (*Cbp_Co_*, *Cbp_Cg_*). The significance of difference between the subgenome convergence indices is also indicated (binomial test,***, *p* < 0.001). For both graphs, the number of differentially expressed genes considered for each tissue are indicated (N).

In flowers, median *S* values for genes that showed significant differential expression between parental species (*FDR* < 0.05) showed dominance of the *Cbp_Co_* over the *Cbp_Cg_* subgenome (∆_*S*_= 0.07), while the opposite pattern – *i.e.* dominance of *Cbp_Cg_* over *Cbp_Co_* – was observed in leaves and roots (∆_*S*_= −0.08 and −0.14, respectively; Fig. 6A). This pattern was also observed when considering all genes, though it was less pronounced (Fig. S4). Such a dominance cannot only be due to the genes showing strict dominance of one genetic background (*Dominance* category, ~3−5%), but rather indicate a more global dominance of *trans*-regulation of one subgenome. Indeed, even if *S* indices tended to show a large legacy of parental genome expression, positive correlations between *S_Cg_* and *S_Co_* (Spearman’s *ρ*, all *p* < 0.001) confirmed that both subgenomes were co-regulated in the same direction (Fig. S4), towards *C. orientalis* in flower tissues and towards *C. grandiflora* in leaf and root tissues.

Finally, in all tissues, most convergence indices were positive (Fig. 6B and S5), indicating that the difference in gene expression between subgenomes (∆_*sub*_) was generally lower than the difference between parental species (∆_*par*_); also, the larger the difference in expression between parental species, ∆_*par*_, the stronger the convergence between subgenomes, *C_Cbp_* (Spearman’s *ρ* = 0.63*, ρ* = 0.74*, ρ* = 0.66, respectively for flowers, leaves and roots; all *p* < 0.001). However, although the overall degree of convergence was the same in the three tissues, the convergence was not symmetrical between the two subgenomes. In flowers, *Cbp_Cg_* tended to shift more towards *Cbp_Co_* than the converse (*C_Cg_ > C_Co_*, Fig. 6B), while the opposite was true in the two other tissues (*C_Co_ > C_Cg_*, Fig. 6B). This explains the dominance patterns observed through the *S* indices and confirms the role of unbalanced *trans*-regulation in that system.

### Genes showing converging expression patterns are enriched for specific functions

Regardless of the tissue considered, the expression profiles did not correspond to specific physical clusters along the genome with transcripts belonging to a given profile being spread across the genome: for each scaffold and each category, the average distance (bp) between two transcripts randomly sampled within a given category was not significantly different than that of two transcripts randomly sampled in different categories (Wilcoxon-Mann-Whitney’s test, all *p* > 0.05, Fig. S6). This suggests that the differential expression is not driven by large-scale epigenetic changes along chromosomes.

Gene ontology analyses, revealed, however, that the different expression profile categories (Fig. 5) were enriched for different molecular functions (MF, average overlap between categories: 17, 20 and 19% for flowers, leaves and roots tissues, respectively, Table S6A) and biological processes (average overlap, 10, 7 and 13%, Table S6B), though neither MF nor BP of a given category tended to cluster into specific networks). At the tissue level, the different expression profile categories were enriched for different MF and BP with a small average overlap between tissues (MF, 7% and BP, 7%, Table S7A and B), highlighting the specificity of expression regulation in different tissues.

We showed above that the main difference in expression between tissues was in the convergence of the two subgenomes: in flowers, *Cbp_Cg_* expression pattern converged toward that of *Cbp_Co_*, while for the two other tissues convergence was in the opposite direction (*Cbp_Co_* toward *Cbp_Cg_*). We tested whether the transcripts showing a convergence of *Cbp_Cg_* toward *Cbp_Co_* (hereafter, *Conv_Co_* genes) or a convergence of *Cbp_Co_* toward *Cbp_Cg_* (hereafter, *Conv_Cg_* genes) were enriched for different molecular functions and biological processes. The two gene sets, *Conv_Co_* or *Conv_Cg_* genes, were indeed enriched for GO terms belonging to different clusters (Fig. S7). For instance, in the flower tissues, *Conv_Co_* genes are enriched for biological processes involved in the transition between vegetative and reproductive phases, the dormancy of floral meristems and male meiosis, while *Conv_Cg_* genes were enriched for cell redox homeostasis and related biological processes (Fig. S7 and Fig. S8). As expected, underlying molecular functions also tended to group into distinct clusters corresponding to different functional networks (Fig. S7 and Fig. S8). Finally, the two gene sets were also enriched for the same biological processes (e.g., fatty acid biosynthesis in flowers, sucrose and carbohydrate metabolisms in leaves and general metabolism in roots) or molecular functions (e.g, RNA, nucleotide and GTP binding or MF related to transporter activity, Fig. S7 and Fig. S8) indicating concerted changes of gene expression between the two subgenomes.

### Deleterious mutations accumulate preferentially on the *C. orientalis* subgenome and are associated with the level of expression

Among the 11 million genomic sites segregating across the five genomes, about 3 million alleles were specific to the *Capsella* species, and 669,675 of these species-specific alleles were annotated by SIFT4G with the *A. thaliana* SIFT database, and 432,354 of them were annotated with the *C. rubella* database.

The estimated proportion of deleterious mutations among species and among the four populations of *C. bursa-pastoris* were similar independently of whether *A. thaliana* or *C. rubella* was used for SIFT4G annotation (Fig. 7A and Fig. S9A). Despite a lower number of accessions, the same pattern as in (35) was observed: i) the *C. grandiflora* genome had a lower proportion of deleterious mutations than *C. orientalis* or either subgenome of *C. bursa-pastoris* ii) within *C. bursa-pastoris*, the *Cbp_Cg_* subgenome always had a lower proportion of deleterious mutations than the *Cbp_Co_* subgenome of the same population and iii) among the *C. bursa-pastoris* populations, both subgenomes of the Asian population had a higher proportion of deleterious mutations than the corresponding subgenomes in the other three populations, indicating a higher rate of mutation accumulation in this population. The proportion of deleterious mutations of the newly added CASI population was most similar to that of the EUR population with a larger variance of the proportion of deleterious mutations carried by *Cbp_Cg_* subgenome of CASI accessions (Fig. 7A).

**Fig. 7.**
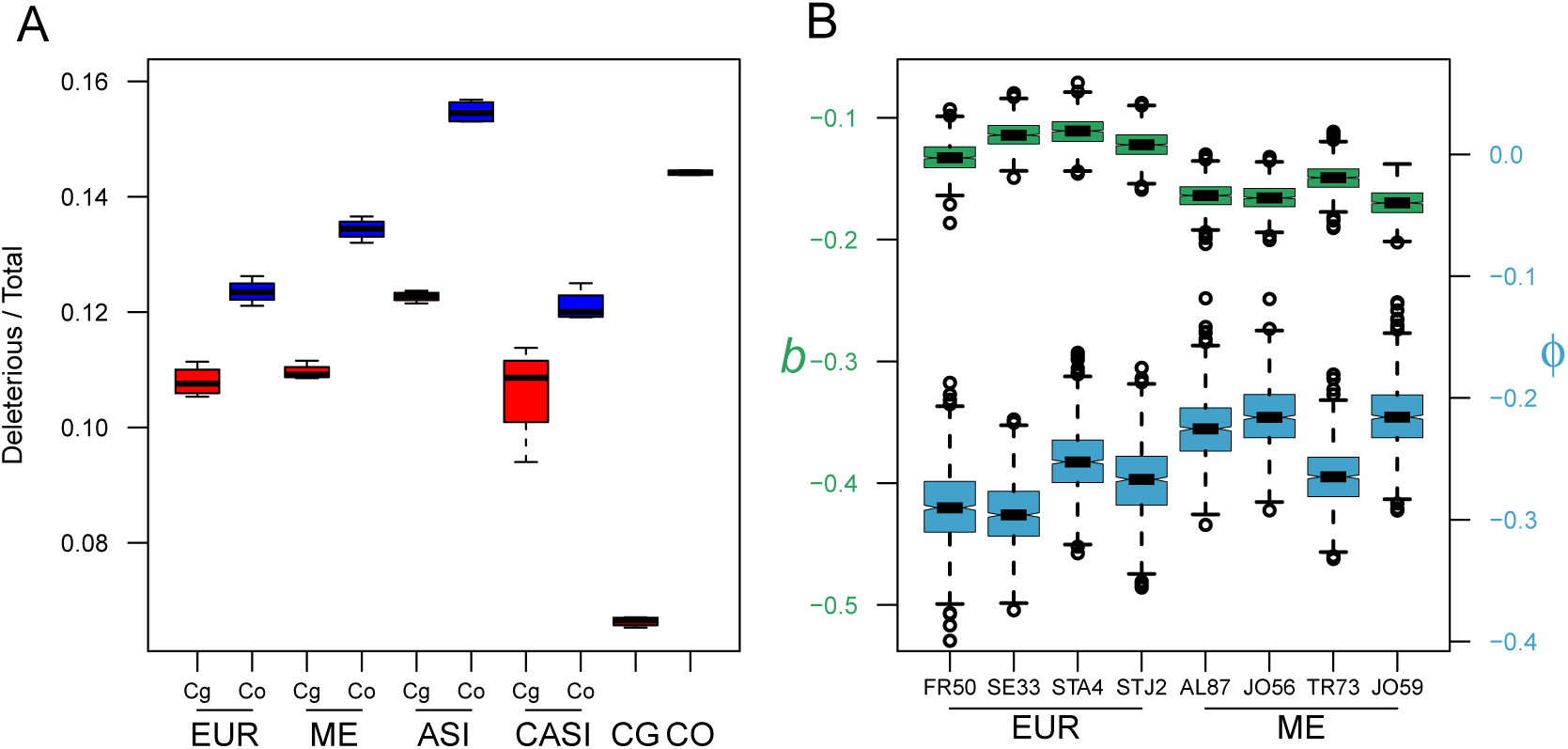
Variation in deleterious mutations in the two subgenomes of *C. bursa-pastoris*. **A.** Proportion of deleterious mutations in the subgenomes and in the parental species. CO, CG, ASI, EUR, ME, CASI correspond to *C. orientalis*, *C. grandiflora*, and four populations of *C. bursa-pastoris*, respectively. The two subgenomes are indicated with Co and Cg. Functional effects were annotated with the *C. rubella* SIFT database (the annotation with *A. thaliana* SIFT database is in the Fig. S9). **B.** Maximum likelihood estimates of parameters of the distribution of deleterious mutations on *Cbp_Cg_* genes. Each box represents the estimates of one accession, with 1000 bootstrap replicates. The estimates are presented as the difference between the estimated parameter for deleterious mutations and the estimated parameter for synonymous mutations (*b*=*b*DEL-*b*SYN, *φ*= *φ*DEL - *φ*SYN). Notches represent the median and the 95% confidence interval. The left axis shows estimates of the bias parameter, *b* (red boxplots), and the right axis shows estimates of the variance parameter *φ* (blue boxplots). The estimated parameters for DEL and SYN are shown separately in Fig. S10.

Mutation accumulation pattern between the two subgenomes was further investigated by estimating the mutation accumulation bias towards *Cbp_Cg_*, *b*, and the overdispersion parameter *φ*. *b* was positive for *SY N* indicating a mapping bias towards *Cbp_Cg_*. *b* was also positive for *DEL* mutations in all accessions (Fig. S10), but much smaller than SYN and therefore *b*DEL *b*SYN was negative (Fig. 7B). This indicates a general bias towards more *DEL* mutations in the *Cbp_Co_* subgenome (*b*DEL *< b*SYN). The same pattern was observed for *φ* (*φ*DEL *< φ*SYN), the difference *φ*DEL *φ*SYN being also negative (Fig. 7B and Fig. S10). Hence, contrary to the expectation of the pseudogenization process, the distribution of deleterious mutations was less over-dispersed than expected at random, suggesting that the accumulation of too many deleterious mutations per gene is prevented, a mechanism that might contribute to the maintenance of both homeologue copies. However, it should be noted that more silenced genes were observed in *Cbp_Co_* than in *Cbp_Cg_*. (Fig. S11). Finally, a significant association between the deleterious mutations bias (*d_DEL_*) and the homeologue expression bias (*e*) was found for all three tissues (Fisher’s exact test, all *p* < 0.001): the categories where deleterious mutations and expression bias varied in the same direction (*i.e.*, *d_DEL_ >* 0 and *e >* 0 or *d_DEL_* and *e* < 0) were over-represented (Table S8). The homeologue copy carrying the highest number of deleterious mutations thus tends to show the lowest expression level. No such association was found when considering only synonymous mutations (Fisher’s exact test, *p* = 0.57, 0.74 and 0.27 for flowers, leaves and roots tissues, respectively), confirming that the association between deleterious mutations and expression level was not the result of a mapping or annotation bias toward one of the two subgenomes (Table S8).

## Discussion

The events accompanying the birth of a polyploid species have often been described in rather dramatic terms, with expressions such as “transcriptomic shock” or “massive genome-wide transcriptomic response” often used (*e.g.* (7, 55, 56)). The early and formative years of a young polyploid might indeed be eventful, but what happens afterward may well be less dramatic, especially for tetraploid species with a disomic inheritance such as the shepherd’s purse. In the present study, we compared some of the genomic and transcriptomic changes that occurred between *C. bursa-pastoris* and its two parental species *C. grandiflora* and *C. orientalis*. Overall, the emerging picture is one of an orderly and rather conservative transition towards a new “normal” state. A conservative transition, because after around 100,000 generations we can still detect a significant parental legacy effect on both the number of deleterious mutations accumulated and gene expression patterns. And an orderly one too, since the emerging pattern of expression involves a balance between *cis*-and *trans*-regulatory changes suggesting the emergence of coordinated functioning of the two subgenomes. This general impression of a non-stochastic transition process to polyploidy (57) is reinforced by the variation in patterns of gene expression across the three tissues: as one would expect, the expression of both subgenomes in selfing *C. bursa-pastoris* was biased towards the selfing parent *C. orientalis* in flower, whereas in leaf expression of the two subgenomes were mostly similar, and in roots expression was biased towards *C. grandiflora*. This expression bias towards the *C. orientalis* subgenome in flowers despite a higher accumulation of deleterious mutations in this subgenome suggest that the evolution of gene expression is not entirely random.

### Demography and expression: a limited effect of introgression?

Previous studies have stressed the importance of population structure and demographic history in genomic and transcriptomic studies of *C. bursa-pastoris* (34, 35); (35), for instance, showed a significant introgression of *C. orientalis* genetic background into Asian populations of *C. bursapastoris*. In the present study, we indeed showed that overall gene expression pattern reflected the main phylogenetic relationships. Each subgenome was the closest to the parental species it was inherited from and populations from close ge-ographic areas tended to cluster together, except for Central Asian accessions (CASI), which clustered with European ones even if they were geographically closer to the Asian or Middle-East ones. Most likely these samples were recently introduced to Central Asia, as it was suggested for *C. burs pastoris* accessions with European ancestry inhabiting the Russian Far East (33).

When comparing the number of differentially expressed genes between *C. bursa-pastoris* and parental species, no specific trend was detected and Asian accessions were not the closest to *C. orientalis* as one would have expected because of introgression. In leaf and roots tissues ASI was even closer to *C. grandiflora* than to *C. orientalis*. This can be explained by the fact that the vast majority of the genes (up to 80%) did not show any difference in expression (thus hiding a more subtle signal). Assessing the influence of introgression on expression pattern would require a more thorough investigation, for instance by focusing on genes for which introgression was actually characterized.

### Transition to polyploidy: compensatory *cis*-*trans*-effects and stabilizing selection

As mentioned above, in the case of a newly formed allopolyploid species one would expect the two copies of a gene to be under the influence of *trans*-regulatory elements inherited from both parents and its expression level to first move towards the mean expression of the two parental species. However, different forces could lead to an excess of divergence in subgenome expression compared to what would be expected under a pure drift model. Polyploidy creates a large redundancy in gene function that should free one of the copies from purifying selection. Generally, the copy carrying more deleterious mutations is expected to degenerate, biasing the expression pattern toward one of the two parental species, even if sub-or neo-functionalization can still occur but to a much lower extent. This ought to be particularly true for *C. bursa-pastoris* as one of its parental species, *C. orientalis*, is a selfer that has accumulated more deleterious mutations than the other parent, the outcrossing *C. grandiflora* (35). This process will be reinforced by the enhancer runways process (25), that should strengthen *cis*-acting elements from the *Cbp_Cg_* subgenome as the *Cbp_Cg_* subgenome has a higher heterozygosity and a lower genetic load than the *Cbp_Co_* subgenome.

In our study, however, we did not observe any “transcriptomic shock” (as for instance in, (7, 55)) neither major homeologue expression remodeling and/or subgenome expression asymmetry (as in *e.g.* (58)). In contrast, our study, like some others before it (15, 57, 59, 60), instead suggests an overall conservation of the expression pattern in polyploids and hybrids. And even if a “transcriptomic shock” did take place during the formation of the tetraploid, expression changes have stabilized since then. Some 100,000 years later parental legacy on subgenome expression is still detectable and the two subgenomes’ expression patterns are still closer to each other than that of parental species, clearly indicating that none of the subgenomes has degenerated; as expected, however, the *Cbp_Co_* subgenome carries more silenced genes and a higher proportion of deleterious mutations than *Cbp_Cg_*. Most of the genes were under both *cis-* and *trans-*acting elements; the *No difference* and *Intermediate* expression categories represented up to 70 to 80% of genes depending on the tissue considered. Only a small fraction (5 to 10%) of genes showed either almost pure *cis-* (*Legacy* category) or *trans-* regulation (*Dominance* category). While the former can be explained by the absence of crossed *trans-*regulation, the latter could be due to the dominance of transcription factor of one subgenome over the other; though, in both cases, post-hybridization mutations affecting either *cis*-or *trans*-acting elements or both could have evolved. The remaining fraction (up to 15%, *Reverse*, *Compensatory-drift* and *Transgressive*) showed a more complex pattern that is hard to assign to a simple factor but could be in part due to new, intertwined *cis-* and *trans-*regulation across subgenomes. It should be noted that such patterns can naturally emerge after hybridization as a byproduct of stabilizing selection on diverging optima (61) for *Transgressive* profiles, on the overall amount of protein produced for *Compensatory-drift* profile, and on intermediate level of expression for *Reverse* profile, without invoking additional specific processes. To address further this question, it would be interesting to compare auto-and allopolyploids to tease apart the effects of hybridization and genome doubling.

Even though this does not, in any way, alter the conclusion above, we also would like to note here that the classification of overall expression patterns used in Table 1 and 2 in different categories is somewhat arbitrary as some expression patterns are ambiguous and could have been classified in different categories. It should also be pointed out that these classifications were dependent on the chosen False Discovery Rate (FDR). As a control, we reproduced the analysis based on unphased data of *Cbp* expression, with FDR< 0.01 and 0.1 (Table S9). It indicated that the number of genes within the different categories can vary substantially with the different FDR level (mainly because of variation in *No difference* category), however, the main patterns were not altered. Moreover, the main pattern of variation we described was a change in dominance between tissue that is obviously not affected by the bias described before. In part to overcome the limitations inherent to any *a priori* classification, we developed the expression similarity index, *S*, that confirmed our conclusions.

### Level of expression dominance varies across tissues and functions

Allopolyploid species are often examined for unequal expression between homeologous genes because of their hybrid nature but other aspects of gene expression have been less extensively studied. For example, there might be no difference in the relative expression of subgenomes (balanced homeologue expression), but the total amount of transcripts can vary and reflect the dominance of the level of expression of one of the parents (62). *C. bursa-pastoris* exhibits rather balanced homeologue expression, but the summed expression of the two homeologues shows differentiation across tissues with the dominance of *C. orientalis* expression level in flowers, and *C. grandiflora* level in leaves and roots. The genes with significant expression bias between subgenomes also show strong dominance of *Cbp_Co_* expression over *Cbp_Cg_* in flower. However, a positive correlation between the expression deviation indices of the two subgenomes indicates that this dominance is not primarily caused by up-regulation or down-regulation of one parental copy, but rather unidirectional regulation of homeologous genes as it has been observed, for instance, in cotton and coffee (2, 30, 63). This convergence could be possible because of the low divergence between the subgenomes of *C. bursapastoris* and, hence, the absence of barriers for *trans*-acting regulation of homeologous genes.

An intuitive explanation of this bias in flower tissues could be that this simply reflects the fact that both *C. orientalis* and *C. bursa-pastoris* are selfing species with tiny flowers, in contrast to *C. grandiflora*, an outcrossing species that has large flowers. A way to test this hypothesis would be to compare *C. orientalis* with both *C. grandiflora* and *C. rubella* for the genes implicated in the bias towards *C. orientalis* using root tissues as a control. In contrast, in the non-reproductive leaf and root tissues, expression is biased towards the genome of the outcrossing *C. grandiflora*. Although this interpretation needs further validation, it stands against the genomic shock pattern that implies a disruption of expression patterns. Finally, although the bias of expression observed between homeologous genes is not strongly shifted towards either subgenome, it is not random either: one subgenome can dominate over the other for a given function or pathway in a given tissue, suggesting constrained evolution in gene expression regulation at a tissue/function level. In many cases, it is not straightforward to explain why a particular subgenome dominates for a particular function, and this could simply be the result of coincidence in neutral evolution of gene regulation networks. In other cases such as flower tissues, however, the observed dominance makes biological sense.

### Both subgenomes of *C. bursa-pastoris* are maintained, but they are not equal

Redundancy of polyploid genomes often assumes evolution of non-functionalization of duplicated genes (64–66) or even of a whole subgenome (67–69). When one gene copy of a duplicated gene starts to degenerate, the purifying selection on that copy becomes weaker and the deleterious mutations accumulate further, while the other copy of the gene remains functional and under purifying selection. If non-functionalization is prevalent, deleterious mutations are expected to be more unevenly distributed between the homeologous genes and even between the two subgenomes. We indeed observed more deleterious load in the *Cbp_Co_* subgenome with the absolute load comparison and with the estimated parameter *b* indicating its degeneration. However, the dispersion for deleterious mutations indicated that they tend to be more evenly distributed between the homeologous genes than expected at random. This suggests that *Cbp_Co_* genes cannot degenerate further after a certain amount of genetic load is accumulated. Thus, although the amount of accumulated genetic load differs between subgenomes of *C. bursa-pastoris*, both subgenomes are maintained and there is no large-scale non-functionalization at the gene and subgenome levels.

One might expect the differences between homeologues in accumulation of deleterious mutations would lead to bias in gene expression. For example, *Arabidopsis suecica*, like *C. bursa-pastoris*, is an allopolyploid species with parents characterized by different mating systems: the outcrossing *Arabidopsis arenosa*, and the selfing *Arabidopsis thaliana* (70). Chang *et al.* (71) observed a bias in expression in favor of the *A. arenosa* subgenome and, among other hypotheses, suggested that this bias could be due to the fact that mildly deleterious alleles are not purged as efficiently from the *A. thaliana* subgenome as from the *A. arenosa* subgenome. In *C. bursa-pastoris*, the *Cbp_Co_* subgenome had a higher proportion of deleterious mutations than the *Cbp_Cg_* subgenome, but there was no strong bias in expression between subgenomes. However, when we paired the amount of derived deleterious mutations with the expression level of each gene and compared homeologous genes, we found that there was a significant association between deleterious mutation bias and expression bias (Table S8). The homeologous gene with more deleterious mutations tends to have a lower expression level than the other one. Moreover, we also found that there are more silenced genes in *Cbp_Co_*, which is the subgenome with a higher proportion of deleterious mutations. These results are in accordance with the hypothesis that the bias in expression is linked to the accumulation of deleterious mutations. Yet, it is worth noting that the expression bias may not necessarily be the result of the biased distribution of deleterious mutations. The homeologue expression bias could also be the cause of the observed deleterious mutation bias, especially considering that we have only investigated the deleterious mutations in coding regions. Purifying selection on the homeologue with lower expression can be weaker (72), therefore it is less efficient in eliminating deleterious mutations. At any rate, the fact that we have a relative dominance of expression of *Cbp_Co_* in flowers and of *Cbp_Cg_* in other tissues, despite *Cbp_Co_* subgenome having a higher proportion of deleterious mutations than *Cbp_Cg_*, suggests that parental legacy and functional constraints may also play a major role.

## Conclusion

In 1929, George Shull, one of the most prominent geneticists of his time (73), wrote: “It is considered a matter of fundamental significance that the increase in a number of chromosomes in the *bursa-pastoris* group is correlated with greater variability, greater adaptability, greater vigor, and greater hardiness”. In the present study, the merging of the two parental genomes was not accompanied by major disruptions of the transcriptome. Instead, there was a strong parental legacy and the emergence of a shift in the subgenome expression pattern towards a new “equilibrium” state reflecting the composite nature of the new species. Hence, being a selfer like its *C. orientalis* parent, there was a shift in flower tissues of the expression pattern of the *C. grandiflora* subgenome towards that of *C. orientalis*. Similarly, it seems also possible that the dominance of the *C. grandiflora* inherited subgenome in roots and leaves contributed to the high competitive ability of *C. bursa-pastoris*, which was similar to that of *C. grandiflora* but much higher than that of *C. orientalis* and *C. rubella*, its two self-fertilizing congeners (74, 75). It therefore seems that the present study, together with those more focused on fitness of *C. bursa-pastoris* (74, 75) contributed to better understanding of the causes of the correlation pointed out almost 100 years ago by Shull.

## ACKNOWLEDGEMENTS

We thank Karl Holm, Kerstin Dalman, and Kerstin Jeppsson for help in the lab. Most of the analyses were carried out at Uppsala Multidisciplinary Center for Advanced Computational Science (UPPMAX) under the project b2016212. This study was supported by grants from the Swedish Research Council and the Philip Sörensens Foundation to ML.

## AUTHOR CONTRIBUTIONS

ML, SG, and DK planned the study. DK obtained the data. DK and PM analyzed the genomic data. DK, PM, and MO analyzed the gene expression data. TD performed the genetic load analyses. DK, PM, TD, and ML wrote the article with inputs from all authors. SG, SIW, and ML supervised the project.

---

**Table S1.** Samples information

| Accession | Population | Latitude | Longitude | DNA lib.size | Flower RNA lib.size | Leaf RNA lib.size | Root RNA lib.size | Source (NCBI SRA) |
| --- | --- | --- | --- | --- | --- | --- | --- | --- |
| DL174 | ASI | 38.56 | 121.35 | 76005581 | 74101388 | 65425962 | 51624022 | SRS2762453 |
| JZH152 | ASI | 30.20 | 112.06 | 80652085 | 61417236 | 61334938 | 37507282 | SRS2762444 |
| NJ219 | ASI | 32.03 | 118.46 | 67776611 | 62976278 | 87650066 | 46483576 | SRS2762448 |
| TY118 | ASI | 37.55 | 112.32 | 68816910 | 53541728 | 60561340 | 58656088 | SRS2762440 |
| DUB-RUS9 | CASI | 51.53 | 58.85 | 84128727 | 78319556 | 68099236 | 54375028 | this study |
| KYRG-3-14 | CASI | 39.79 | 72.18 | 45154567 | 66059762 | 60694436 | 57074026 | this study |
| LAB-RUS-4 | CASI | 66.65 | 66.40 | 33843140 | 69017208 | 61974690 | 39439188 | this study |
| TACH-CHIN14 | CASI | 47.07 | 83.01 | 83324082 | 61379494 | 77626154 | 38668184 | this study |
| 85.3 | CG | 39.56 | 20.92 | 57969763 | 55106794 | 68729596 | 59193184 | this study |
| 86.12 | CG | 39.52 | 20.97 | 71852215 | 70392238 | 54187192 | 62579512 | this study |
| 87.26 | CG | 39.48 | 20.98 | 60716530 | 62728886 | 59460660 | 64762960 | this study |
| 88.5 | CG | 39.88 | 20.75 | 45369106 | 67964672 | 44978638 | 47326734 | this study |
| GUB-RUS5 | CO | 51.29 | 58.18 | 55352931 | 58041160 | 55549182 | 61419992 | this study |
| PAR-RUS | CO | 53.30 | 60.10 | 49028787 | 68030766 | 55527408 | 57415936 | this study |
| QH-CHIN4 | CO | 46.70 | 90.83 | 46773214 | 66241216 | 55334438 | 57763206 | this study |
| URAL-RUS4 | CO | 55.11 | 61.39 | 45387760 | 76656450 | 57886342 | 69234050 | this study |
| FR50 | EUR | 48.08 | 7.37 | 70612809 | 57720098 | 46159028 | 33748500 | SRS2762459 |
| SE33 | EUR | 56.15 | 13.77 | 78120829 | 61139124 | 66478952 | 49599858 | SRS2762447 |
| STA4 | EUR | 56.20 | -2.47 | 76063405 | 64453046 | 52318632 | 42635412 | SRS2762460 |
| STJ2 | EUR | 44.51 | -1.21 | 77687488 | 85809658 | 48134448 | 35801122 | SRS2762442 |
| AL87 | ME | 35.45 | 7.96 | 59970536 | 76995752 | 57011620 | 54660030 | SRS2762455 |
| JO56 | ME | 31.97 | 35.98 | 79470603 | 56584612 | 61885264 | 60461952 | SRS2762457 |
| TR73 | ME | 41.02 | 28.97 | 77624846 | 59569756 | 69614876 | 44957844 | SRS2762445 |
| JO59 | ME | 31.97 | 35.98 | 67354447 | 67068132 | 68539844 | 53689276 | this study |

CO, CG, ASI, EUR, ME, CASI correspond to *C. orientalis*, *C. grandiflora*, and four populations of *C. bursa-pastoris*, respectively.

**Table S2.** Differential gene expression between three *Capsella* species in three tissues.

| Tissue | Comparison |  |  |
| --- | --- | --- | --- |
|  | CO vs CG | Cbp vs CG | Cbp vs CO |
| Flowers | 4793 | 4214 | **2796** |
| Leaves | 1729 | **1172** | 1535 |
| Roots | 2742 | **2047** | 3537 |

CO, CG, and Cbp correspond to *C. orientalis*, *C. grandiflora*, and *C. bursa-pastoris*, respectively. The analysis was performed on the unphased expression data of 16,032 genes with the significance level set to 0.05. The smallest differences between species per tissue are in bold.

**Table S3.** Differential gene expression between *Capsella* species/population in three tissues.

| Tissue | Comparison |  |  |  |  |  |  |  |  |  |  |  |  |  |
| --- | --- | --- | --- | --- | --- | --- | --- | --- | --- | --- | --- | --- | --- | --- |
|  | CG vs ASI | CG vs EUR | CG vs ME | CG vs CASI | CO vs ASI | CO vs EUR | CO vs ME | CO vs CASI | ASI vs EUR | ASI vs ME | EUR vs ME | ASI vs CASI | EUR vs CASI | ME vs CASI |
| Flowers | 3946 | 3677 | 3723 | **3395** | 2467 | 3069 | 2358 | **2084** | 907 | 908 | 680 | 645 | **62** | 458 |
| Leaves | **851** | 1192 | 1031 | 1055 | 1154 | 1392 | **978** | 1612 | 394 | 551 | 245 | 360 | **7** | 212 |
| Roots | 1868 | 2123 | **1533** | 1592 | 2167 | 4000 | **2138** | 3305 | 847 | 1052 | 514 | 570 | **7** | 358 |

CO, CG, ASI, CASI, EUR, and ME correspond to *C. orientalis*, *C. grandiflora*, and four populations of *C. bursa-pastoris*, respectively. The analysis was performed on the unphased expression data of 16,032 genes with the significance level set to 0.05. The smallest differences between species/population per tissue are in bold.

**Table S4.**
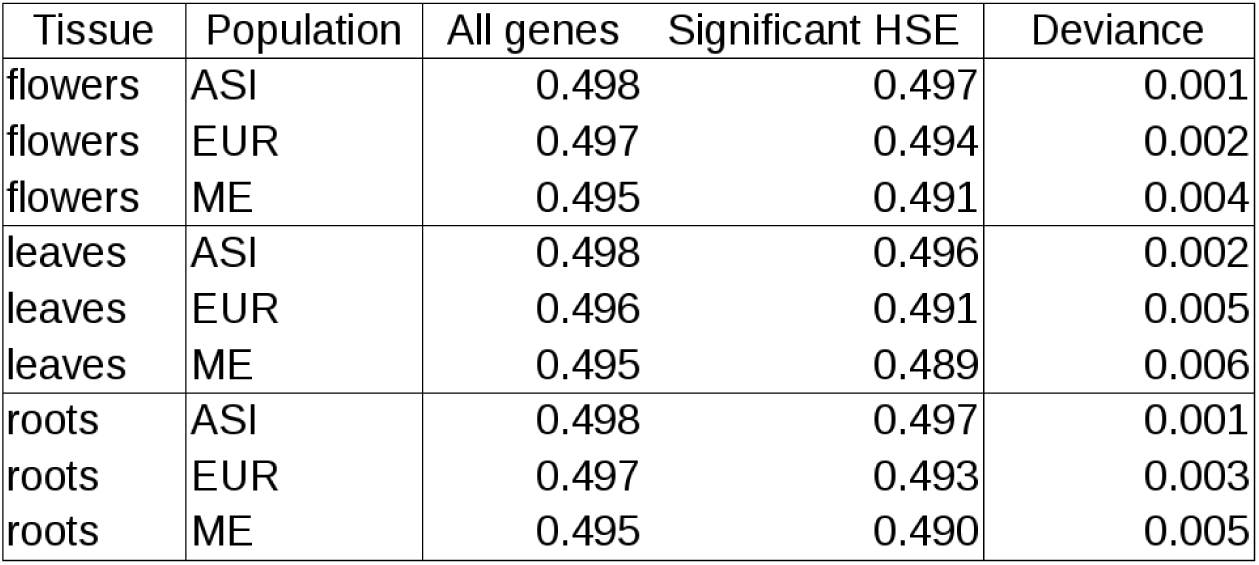
Expression ratio between the two subgenomes of *C. bursa-pastoris* across populations in three tissues. The expression ratio is estimated by the proportion of the *Cbp_Co_* subgenome counts in the total expression counts. The table shows ratios for all assayed genes (All genes), genes with significant homeologue-specific expression (Significant HSE), over Asian (ASI), European (EUR) and Middle Eastern (ME) populations of *C. bursa-pastoris* in three different tissues. The deviance shows the difference in mean expression ration between all genes and genes showing significant HSE.

| Tissue | Population | All genes | Significant HSE | Deviance |
| --- | --- | --- | --- | --- |
| flowers | ASI | 0.498 | 0.497 | 0.001 |
| flowers | EUR | 0.497 | 0.494 | 0.002 |
| flowers | ME | 0.495 | 0.491 | 0.004 |
| leaves | ASI | 0.498 | 0.496 | 0.002 |
| leaves | EUR | 0.496 | 0.491 | 0.005 |
| leaves | ME | 0.495 | 0.489 | 0.006 |
| roots | ASI | 0.498 | 0.497 | 0.001 |
| roots | EUR | 0.497 | 0.493 | 0.003 |
| roots | ME | 0.495 | 0.490 | 0.005 |

**Table S5.** Differentially expressed genes between tissues (A) and populations within tissues (B) for each *C. bursa-pastoris*

| Contrast | *Cbp<sub>Co</sub>* | % | *Cbp<sub>Cg</sub>* | % | Δ% |
| --- | --- | --- | --- | --- | --- |
| **F vs L** | **6474** | **54** | **6475** | **54** | **0** |
| F vs R | 5922 | 50 | 6003 | 50 | -1 |
| L vs R | 5545 | 46 | 5688 | 48 | -1 |

**B**
| Flowers: 4029 |  |  |  |  |  |
| --- | --- | --- | --- | --- | --- |
| Contrast | *Cbp<sub>Co</sub>* | % | *Cbp<sub>Cg</sub>* | % | Δ% |
| ME vs ASI | 279 | 7 | 223 | 6 | 1 |
| ME vs CASI | 214 | 5 | 182 | 5 | 1 |
| ME vs EUR | 216 | 5 | 182 | 5 | 1 |
| ASI vs CASI | 192 | 5 | 198 | 5 | 0 |
| ASI vs EUR | 268 | 7 | 259 | 6 | 0 |
| CASI vs EUR | 88 | 2 | 112 | 3 | -1 |
| Leaves: 3051 |  |  |  |  |  |
| Contrast | *Cbp<sub>Co</sub>* | % | *Cbp<sub>Cg</sub>* | % | Δ% |
| ME vs ASI | 248 | 8 | 236 | 8 | 0 |
| ME vs CASI | 162 | 5 | 134 | 4 | 1 |
| ME vs EUR | 132 | 4 | 133 | 4 | 0 |
| ASI vs CASI | 170 | 6 | 177 | 6 | 0 |
| ASI vs EUR | 156 | 5 | 168 | 6 | 0 |
| CASI vs EUR | 14 | 0 | 20 | 1 | 0 |
| Roots: 3759 |  |  |  |  |  |
| Contrast | *Cbp<sub>Co</sub>* | % | *Cbp<sub>Cg</sub>* | % | Δ% |
| ME vs ASI | 256 | 7 | 243 | 6 | 0 |
| ME vs CASI | 137 | 4 | 125 | 3 | 0 |
| ME vs EUR | 212 | 6 | 180 | 5 | 1 |
| ASI vs CASI | 165 | 4 | 174 | 5 | 0 |
| ASI vs EUR | 251 | 7 | 241 | 6 | 0 |
| CASI vs EUR | 23 | 1 | 34 | 1 | 0 |

ASI: Asia, CASI: central Asia, EUR: Europe and ME: Middle-East. *FDR* < 0.05.

**Table S6.** Overlap between expression profiles in gene ontology term enrichment for Biological processes (A) and Molecular functions (B).

| Biological processes Flowers |  |  |  |  |  |  |  |
| --- | --- | --- | --- | --- | --- | --- | --- |
| Categories | Comp Drift | Dom. CG | Dom. CO | Int. | Leg. | Rev. | Trans. |
| Dominance CG | 10 | - | - | - | - | - | - |
| Dominance CO | 18 | 20 | - | - | - | - | - |
| Intermeate | 16 | 0 | 0 | - | - | - | - |
| Legacy | 12 | 5 | 9 | 19 | - | - | - |
| Reverse | 0 | 0 | 0 | 33 | 0 | - | - |
| Transgressive | 12 | 5 | 5 | 6 | 15 | 17 | - |
| Equal | 15 | 15 | 12 | 16 | 21 | 17 | 21 |

| Biological porcesses Leaves |  |  |  |  |  |  |  |
| --- | --- | --- | --- | --- | --- | --- | --- |
| Categories | Comp Drift | Dom. CG | Dom. CO | Int. | Leg. | Rev. | Trans. |
| Dominance CG | 29 | - | - | - | - | - | - |
| Dominance CO | 0 | 0 | - | - | - | - | - |
| Intermediate | 21 | 10 | 0 | - | - | - | - |
| Legacy | 21 | 5 | 5 | 20 | - | - | - |
| Reverse | 0 | 0 | 0 | 0 | 0 | - | - |
| Transgressive | 0 | 0 | 10 | 0 | 9 | 0 | - |
| No difference | 21 | 11 | 15 | 20 | 9 | 0 | 25 |

| Biological porcesses Roots |  |  |  |  |  |  |  |
| --- | --- | --- | --- | --- | --- | --- | --- |
| Categories | Comp Drift | Dom. CG | Dom. CO | Int. | Leg. | Rev. | Trans. |
| Dominance CG | 26 | - | - | - | - | - | - |
| Dominance CO | 0 | 22 | - | - | - | - | - |
| Intermediate | 5 | 5 | 11 | - | - | - | - |
| Legacy | 14 | 14 | 6 | 10 | - | - | - |
| Reverse | 0 | 0 | 0 | 20 | 20 | - | - |
| Transgressive | 9 | 8 | 22 | 20 | 18 | 20 | - |
| No difference | 26 | 12 | 11 | 20 | 9 | 20 | 22 |

**B**
| Molecular functions Flowers |  |  |  |  |  |  |  |
| --- | --- | --- | --- | --- | --- | --- | --- |
| Categories | Comp Drift | Dom. CG | Dom. CO | Int. | Leg. | Rev. | Trans. |
| Dominance CG | 13 | - | - | - | - | - | - |
| Dominance CO | 11 | 13 | - | - | - | - | - |
| Intermediate | 15 | 27 | 7 | - | - | - | - |
| Legacy | 13 | 27 | 16 | 21 | - | - | - |
| Reverse | 20 | 10 | 30 | 40 | 20 | - | - |
| Transgressive | 22 | 13 | 18 | 17 | 21 | 0 | - |
| No difference | 22 | 7 | 20 | 17 | 29 | 20 | 13 |

| Molecular functions Leaves |  |  |  |  |  |  |  |
| --- | --- | --- | --- | --- | --- | --- | --- |
| Categories | Comp Drift | Dom. CG | Dom. CO | Int. | Leg. | Rev. | Trans. |
| Dominance CG | 12 | - | - | - | - | - | - |
| Dominance CO | 0 | 17 | - | - | - | - | - |
| Intermediate | 20 | 16 | 11 | - | - | - | - |
| Legacy | 14 | 20 | 28 | 20 | - | - | - |
| Reverse | 20 | 40 | 0 | 20 | 20 | - | - |
| Transgressive | 20 | 24 | 28 | 26 | 17 | 60 | - |
| No difference | 20 | 24 | 17 | 9 | 22 | 20 | 14 |

| Molecular functions Roots |  |  |  |  |  |  |  |
| --- | --- | --- | --- | --- | --- | --- | --- |
| Categories | Comp Drift | Dom. CG | Dom. CO | Int. | Leg. | Rev. | Trans. |
| Dominance CG | 3 | - | - | - | - | - | - |
| Dominance CO | 8 | 16 | - | - | - | - | - |
| Intermediate | 17 | 12 | 28 | - | - | - | - |
| Legacy | 20 | 9 | 12 | 18 | - | - | - |
| Reverse | 30 | 10 | 10 | 30 | 10 | - | - |
| Transgressive | 29 | 21 | 12 | 17 | 18 | 60 | - |
| No difference | 17 | 30 | 8 | 25 | 12 | 20 | 13 |

**Table S7.**
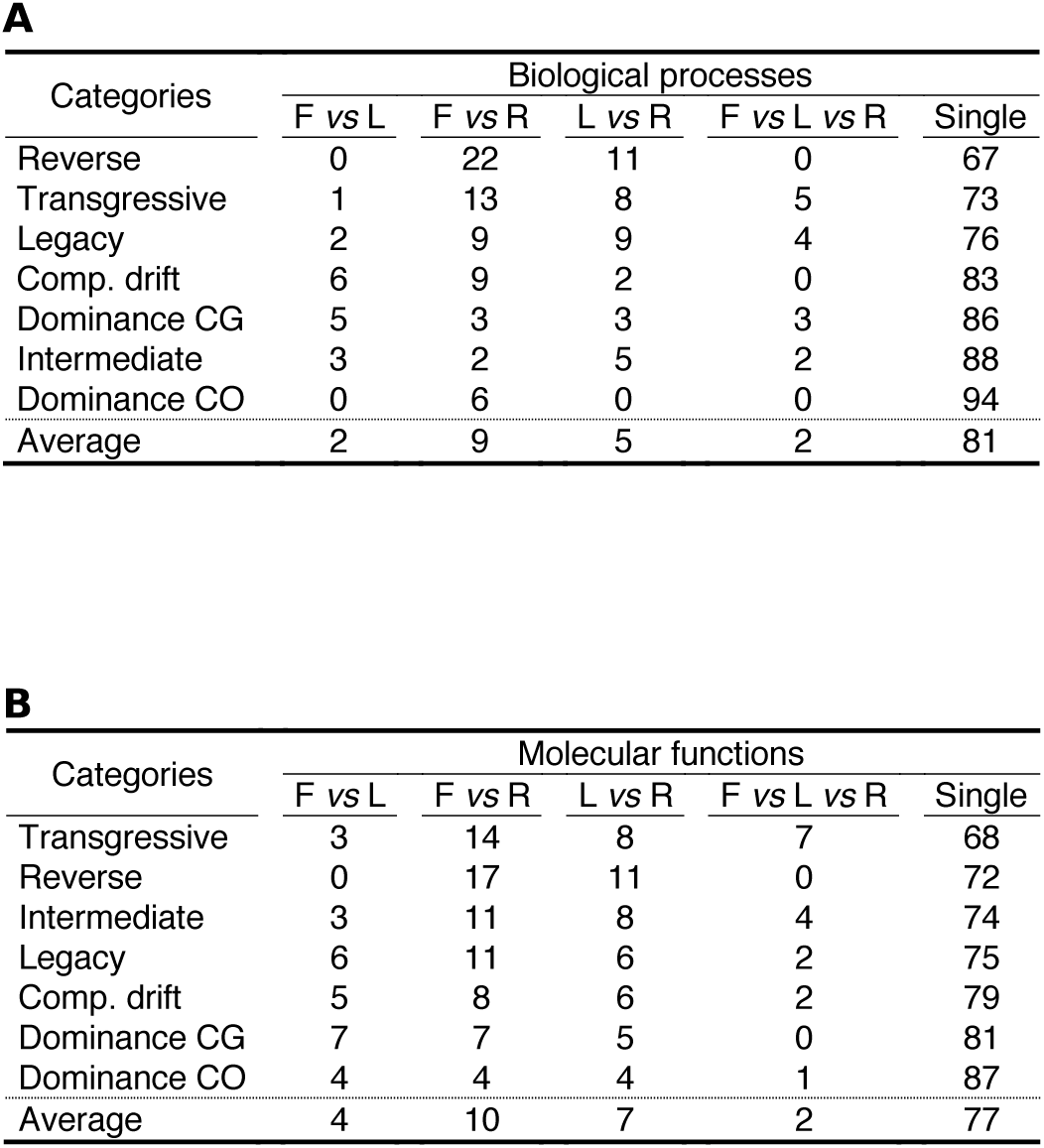
Overlap between tissue in expression profiles gene ontology term enrichment for Biological processes (A) and Molecular functions (B).

**Table S8.** Contingency table of number of genes per category based on deleterious mutation and homelogue expression bias.

| Mutation types | Tissues |  | Obs. (Exp.) |  | Total | p |
| --- | --- | --- | --- | --- | --- | --- |
| | | | $e > 0.5$ | $e < 0.5$ | | |
| DEL | Flowers | $d > 0$ | 5422 (5118) | 5126 (5430) | 10548 | < 2.2E-16 |
| | | $d < 0$ | 2656 (2960) | 3443 (3139) | 6099 | |
|  |  | Total | 8078 | 8569 | 16647 |  |
| | Leaves | $d > 0$ | 5666 (5427) | 5749 (5988) | 11415 | 1.1E-13 |
| | | $d < 0$ | 2862 (3101) | 3660 (3421) | 6522 | |
|  |  | Total | 8528 | 9409 | 17937 |  |
| | Roots | $d > 0$ | 5299 (5128) | 5280 (5451) | 10579 | 2.1E-08 |
| | | $d < 0$ | 2638 (2809) | 3158 (2987) | 5796 | |
|  |  | Total | 7937 | 8438 | 16375 |  |
| SYN | Flowers | $d > 0$ | 9857 (9837) | 10864 (10884) | 20721 | 0.58 |
| | | $d < 0$ | 2907 (2927) | 3258 (3238) | 6165 | |
|  |  | Total | 12764 | 14122 | 26886 |  |
| | Leaves | $d > 0$ | 10189 (10201) | 11432 (11420) | 21621 | 0.74 |
| | | $d < 0$ | 3004 (2992) | 3338 (3350) | 6342 | |
|  |  | Total | 13193 | 14770 | 27963 |  |
| | Roots | $d > 0$ | 9394 (9431) | 10371 (10334) | 19765 | 0.27 |
| | | $d < 0$ | 2848 (2811) | 3042 (3079) | 5890 | |
|  |  | Total | 12242 | 13413 | 25655 |  |

$d$ is the difference of number of mutations (DEL, deleterious and SYN, synonymous) between homeologous copies (DEL<sub>Cg</sub> - DEL<sub>Co</sub> or SYN<sub>Cg</sub> - SYN<sub>Co</sub>) and $e$ is the expression ratio between the two homeologues copies with significant HSE ( $e = \frac{Cbp_{Co}}{Cbp_{Cg} + Cbp_{Co}}$ ). For each category, expected number of gene under category independency hypothesis are given into parenthesis, $p$ is the $p$ -value of Fisher's exact test of independency.

**Table S9.**
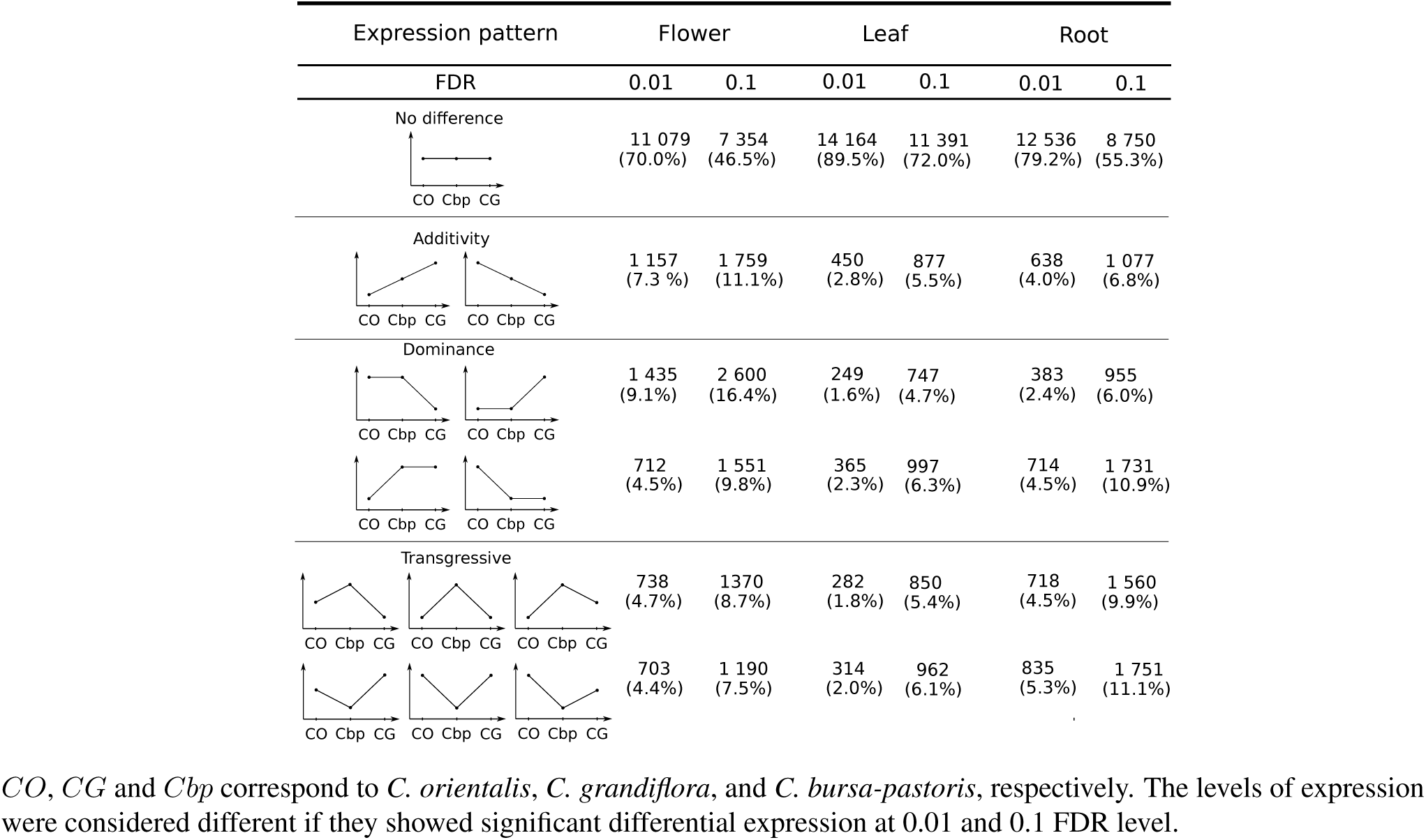
Gene expression levels in *C. bursa-pastoris* and its parental species with different FDR thresholds

| Expression pattern | Flower |  | Leaf |  | Root |  |
| --- | --- | --- | --- | --- | --- | --- |
| FDR | 0.01 | 0.1 | 0.01 | 0.1 | 0.01 | 0.1 |
| **No difference**<br>CO Cbp CG | 11 079<br>(70.0%) | 7 354<br>(46.5%) | 14 164<br>(89.5%) | 11 391<br>(72.0%) | 12 536<br>(79.2%) | 8 750<br>(55.3%) |
| **Additivity**<br>CO Cbp CG CO Cbp CG | 1 157<br>(7.3 %) | 1 759<br>(11.1%) | 450<br>(2.8%) | 877<br>(5.5%) | 638<br>(4.0%) | 1 077<br>(6.8%) |
| **Dominance**<br>CO Cbp CG CO Cbp CG | 1 435<br>(9.1%) | 2 600<br>(16.4%) | 249<br>(1.6%) | 747<br>(4.7%) | 383<br>(2.4%) | 955<br>(6.0%) |
| CO Cbp CG CO Cbp CG | 712<br>(4.5%) | 1 551<br>(9.8%) | 365<br>(2.3%) | 997<br>(6.3%) | 714<br>(4.5%) | 1 731<br>(10.9%) |
| **Transgressive**<br>CO Cbp CG CO Cbp CG CO Cbp CG | 738<br>(4.7%) | 1370<br>(8.7%) | 282<br>(1.8%) | 850<br>(5.4%) | 718<br>(4.5%) | 1 560<br>(9.9%) |
| CO Cbp CG CO Cbp CG CO Cbp CG | 703<br>(4.4%) | 1 190<br>(7.5%) | 314<br>(2.0%) | 962<br>(6.1%) | 835<br>(5.3%) | 1 751<br>(11.1%) |

*CO*, *CG* and *Cbp* correspond to *C. orientalis*, *C. grandiflora*, and *C. bursa-pastoris*, respectively. The levels of expression were considered different if they showed significant differential expression at 0.01 and 0.1 FDR level.

**Fig. S1.**
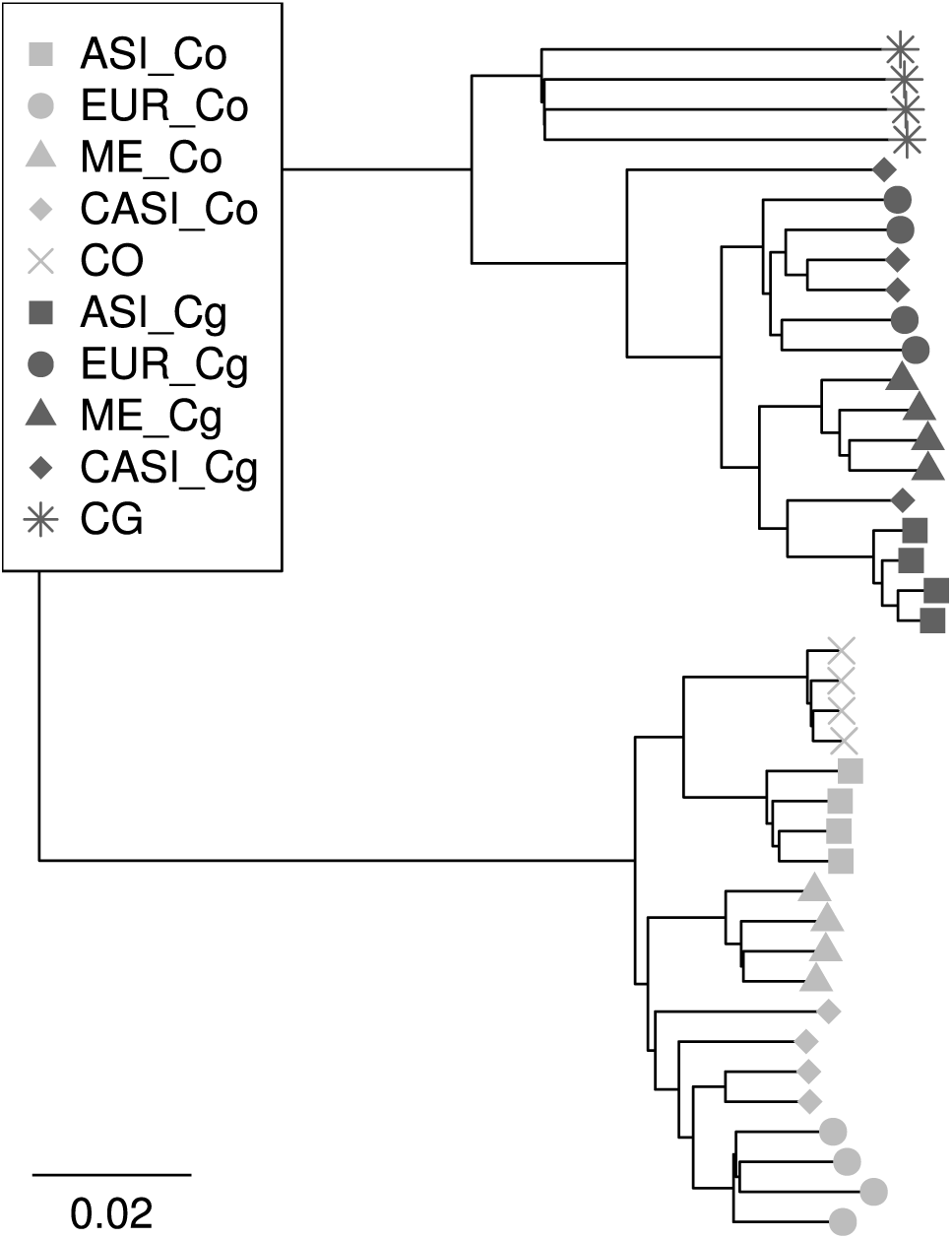
Neighbor-joining tree of the genomic data of three *Capsella species*. CO, CG, ASI, EUR, ME, CASI correspond to textitC. orientalis, *C. grandiflora*, and four populations of *C. bursa-pastoris*, respectively. The two subgenomes are indicated with C_o_ and C_g_. The tree was reconstructed from 11Mb of SNPs and the distance was then scaled to whole genome variation.

**Fig. S2.**
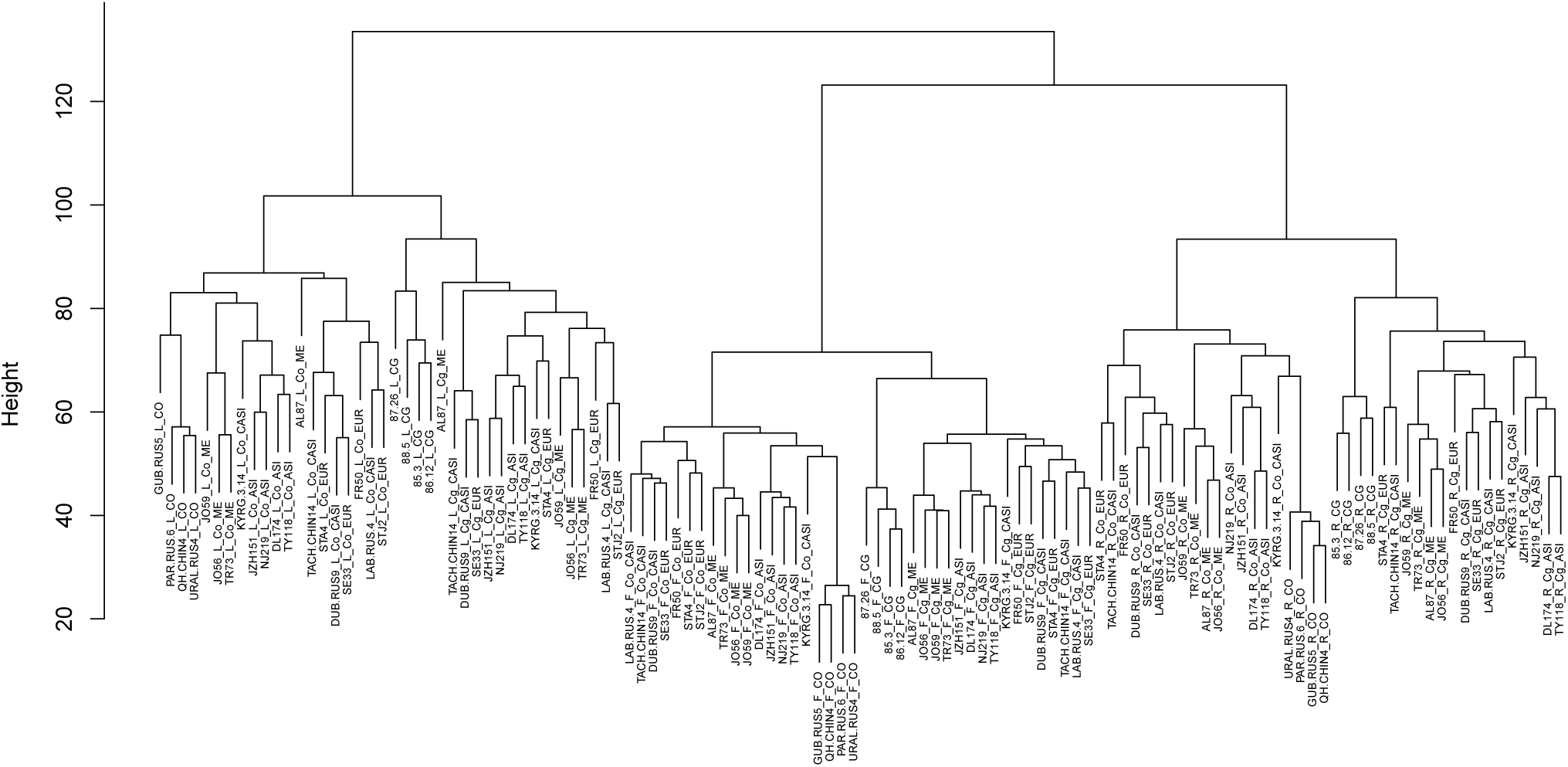
Distance clustering dendrogram of gene expression data of separate samples. Clustering was performed using Euclidean distances and the average agglomerative method on 11959 genes. Each label indicates an accession number, tissue, and population. In the labels, *CO* and *CG* correspond to diploid species *C. orientalis* and *C. grandiflora*, respectively. The Asian, Central Asian, European and Middle Eastern populations of *C. bursa-pastoris* are called ASI, CASI, EUR and ME, and the two subgenomes are indicated with Co and Cg. F, L, and R stand for flower, leave and root tissues, respectively.

**Fig. S3.**
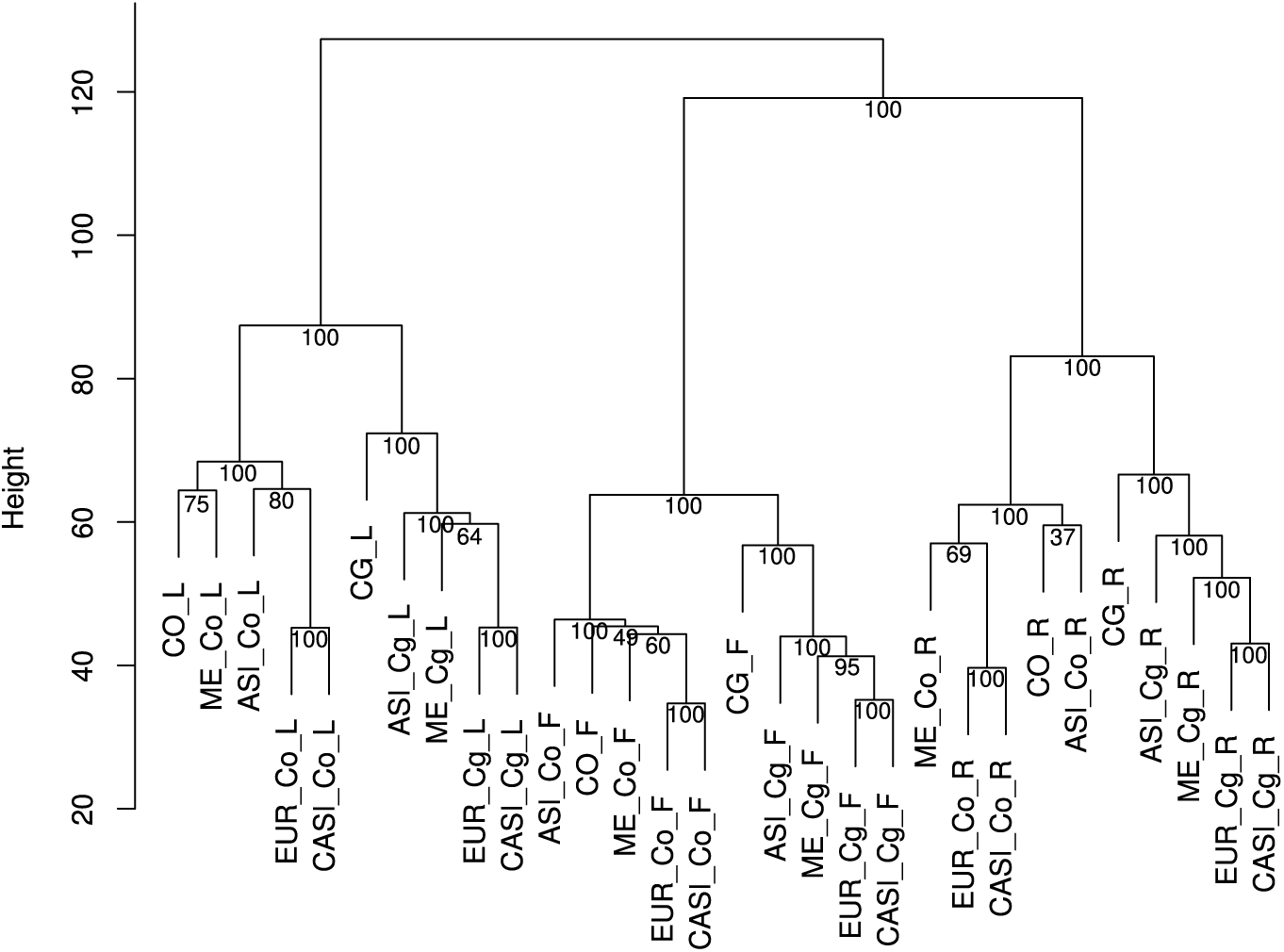
Distance clustering dendrogram of gene expression data. Clustering was performed using Euclidean distances and the average agglomerative method on mean expression values for each population (10,403 genes). *CO* and *CG* correspond to diploid species *C. orientalis* and *C. grandiflora*, respectively. The Asian, European and Middle Eastern populations of *C. bursa-pastoris* are called ASI, EUR and ME, and the two subgenomes are indicated with Co and Cg. F, L, and R stand for flower, leave and root tissues, respectively. Bootstrap support was generated from 1000 replicates.

**Fig. S4.**
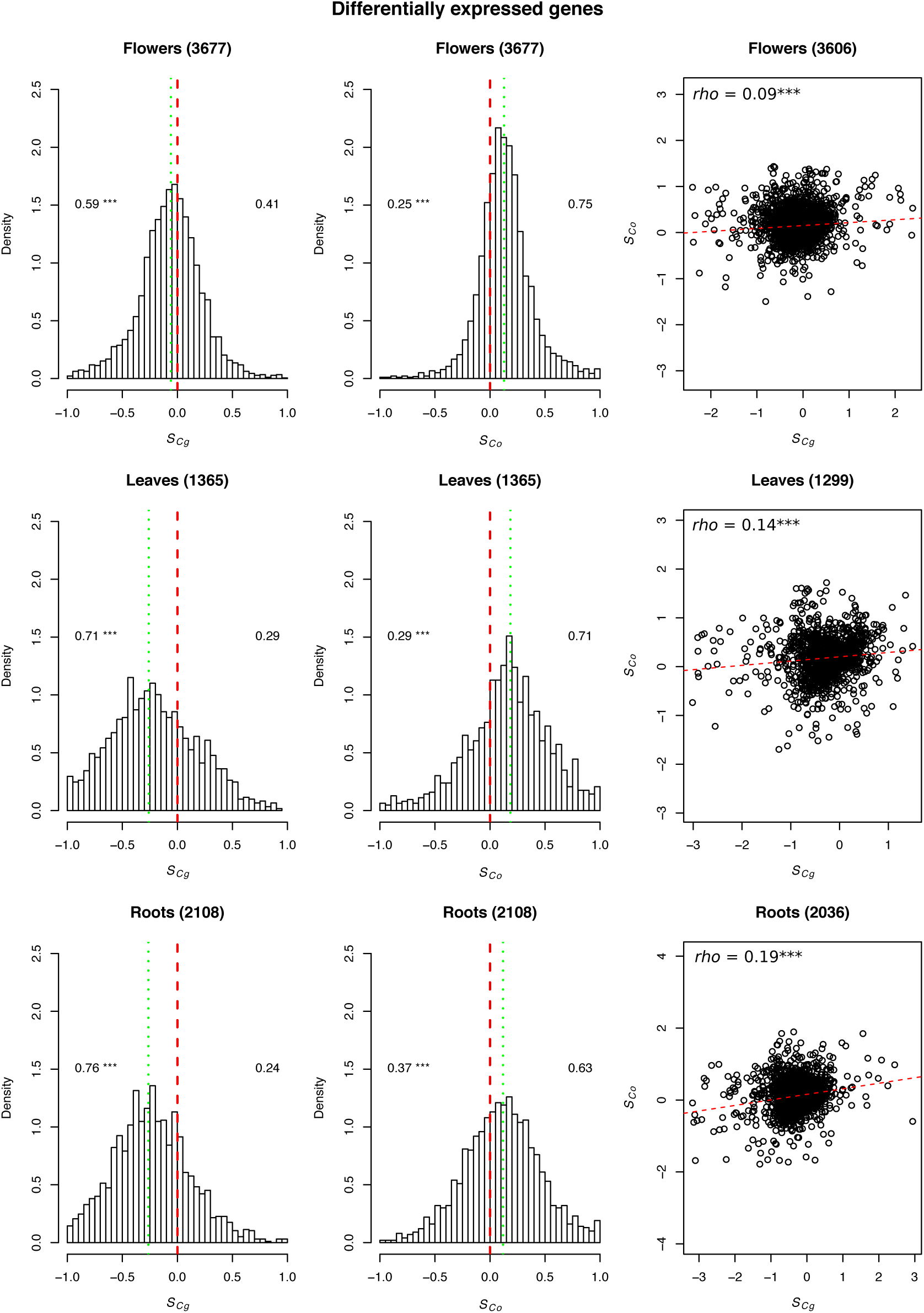
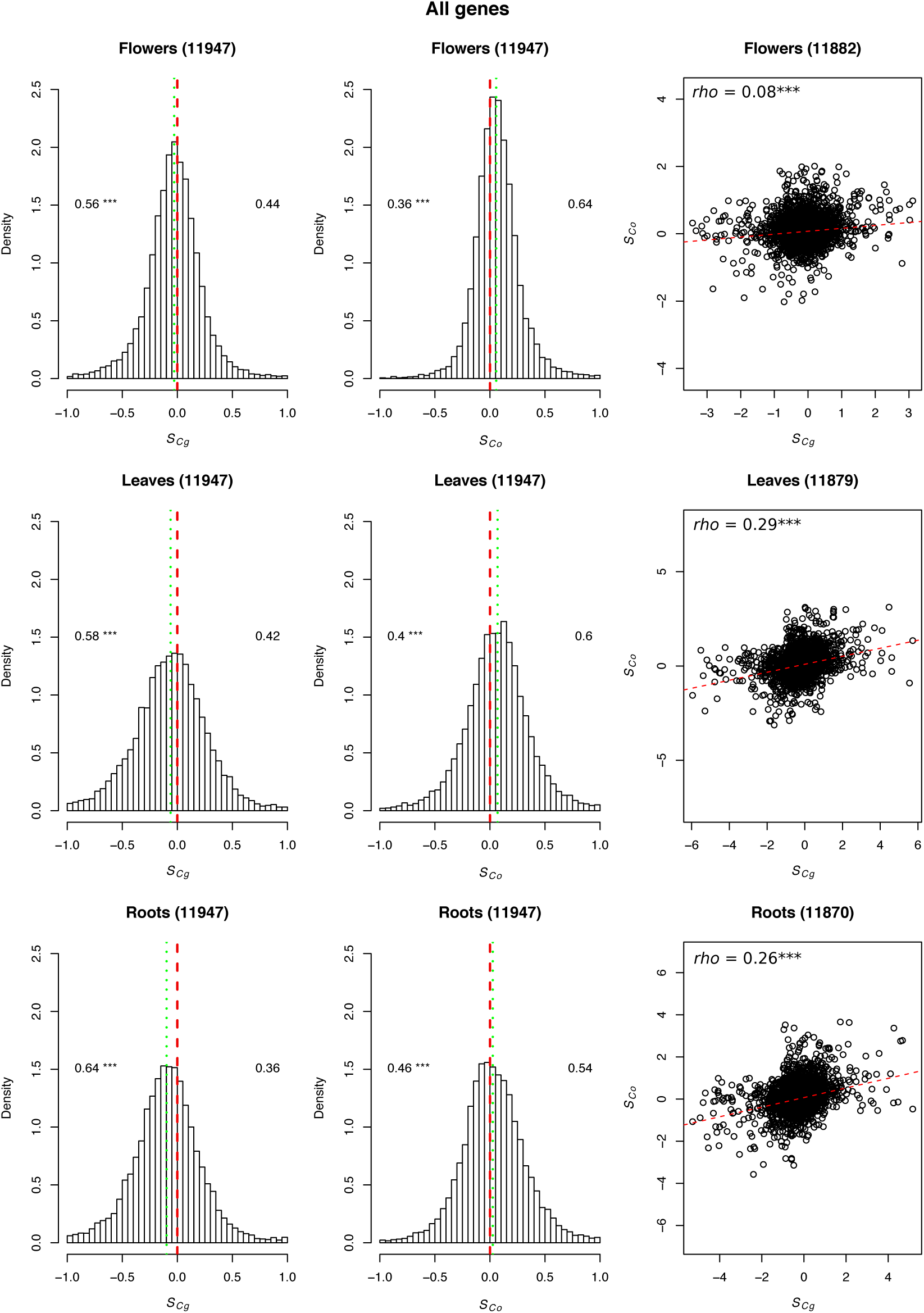
Distribution of the similarity index for each subgenome of *C. bursa-pastoris*. The similarity index was computed for each subgenome (*Cg*, left panels and *Co*, right panels) and for each tissue separately (flowers, top panels, leaves, middle panels and roots bottom panels). The number of genes considered in each analysis is indicated within parentheses. *S* values < 0 means bias towards *CG* while *S* values > 0 bias toward CO. The red dashed line represents *S* = 0 and the green dotted line is the median *S*-value. Numbers are proportions of genes bias toward each parental genome. Note that the x-axis is truncated to the interval [−1,1]. Right panels are correlations between *S_Co_*and *S_Cg_*, red dashed lines are linear regressions between both factors, Spearman’s correlation coefficients (*ρ*) are indicated as well as their significance (***, *p* < 0.001).

**Fig. S5.**
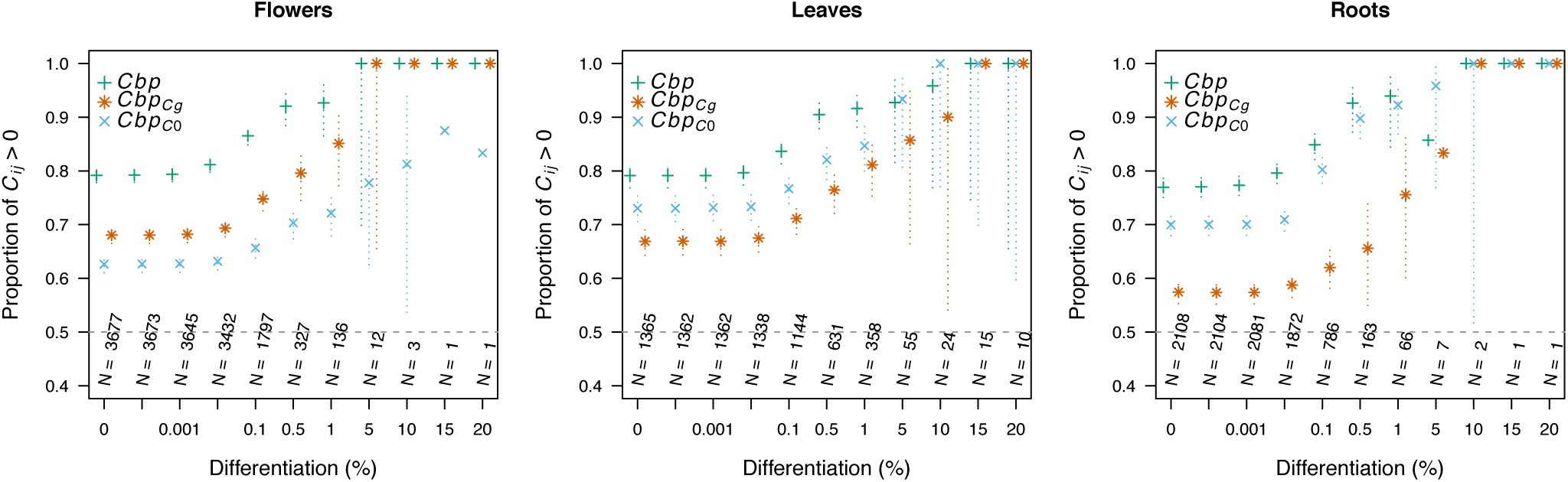
Subgenomes convergence in *C. bursa-pastoris* given differentiation from parental expression. For each tissue, and each subgenome *i*, the proportion of transcripts *j* showing convergence (*i.e.*, *C_ij_>* 0) are reported as a function of differentiation from parental expression with the associated 95% confidence intervals (dotted lines, non-significant when absent). As an indication of power for binomial tests, the number of gene considered for the various threshold of differentiation for the overall *Cbp* convergence are indicated (*N*).

**Fig. S6.**
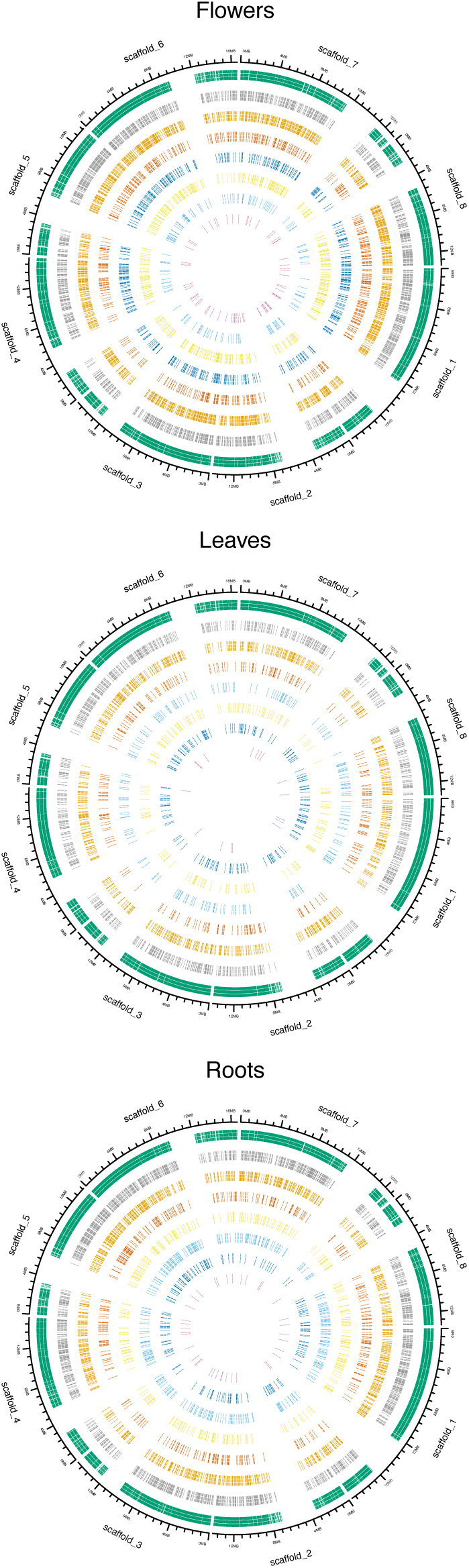
Expression profile regarding genome position. For each tissue, each transcript is positioned on a scaffold by a vertical bar (given *C. rubella* annotation). Concentric circles and colors correspond to the different expression profiles: *No differences*, green; *Transgressive*, grey; *Intermediate*, orange; *Legacy*, red; *Compensatory drift*, yellow; *Dominance*, blue (*Cbp_Co_*dark and *Cbp_Cg_*ligth); *Reverse*, pink. Profiles are organized given their importance (number of transcript).

**Fig. S7.**
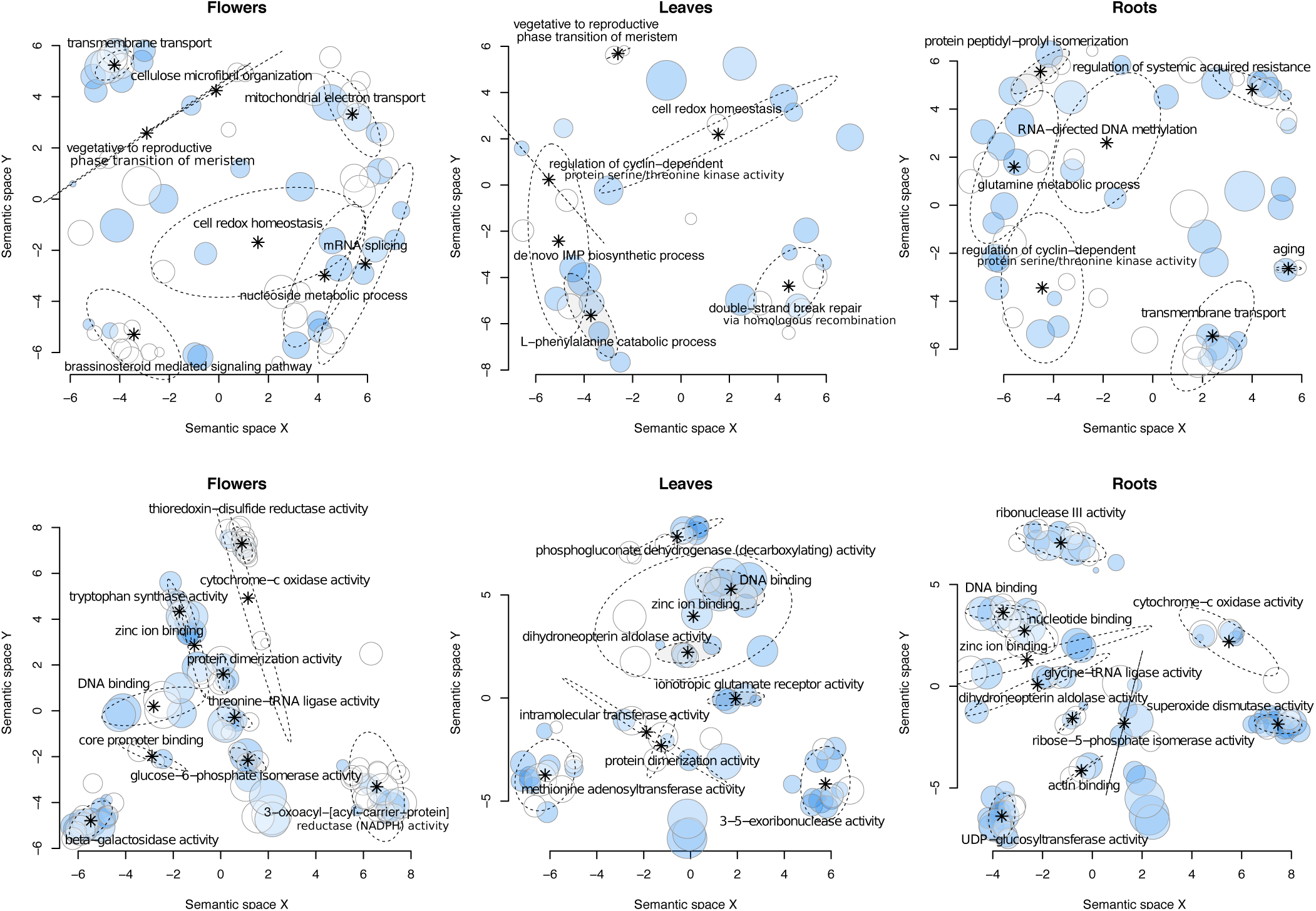
Two-dimensional semantic space representation of significantly enriched GO categories in genes showing convergence in expression. Multidimensional scaling to a matrix of the GO terms’ semantic similarities (see text for details). Semantic representation of GO categories is colored according to the significant over-representation (*FDR* < 0.05) in genes showing a convergence of expression either of *Cbp_Cg_*toward *Cbp_Co_*(white) or of *Cbp_Co_*toward *Cbp_Cg_*(blue) for biological processes (top panels) or molecular functions (bottom panels). The circle diameter is proportional to the number of aggregated GO terms. Ellipses gather GO terms with low redundancy, description of the GO term with the highest dispensability in the group is reported.

**Fig. S8.**
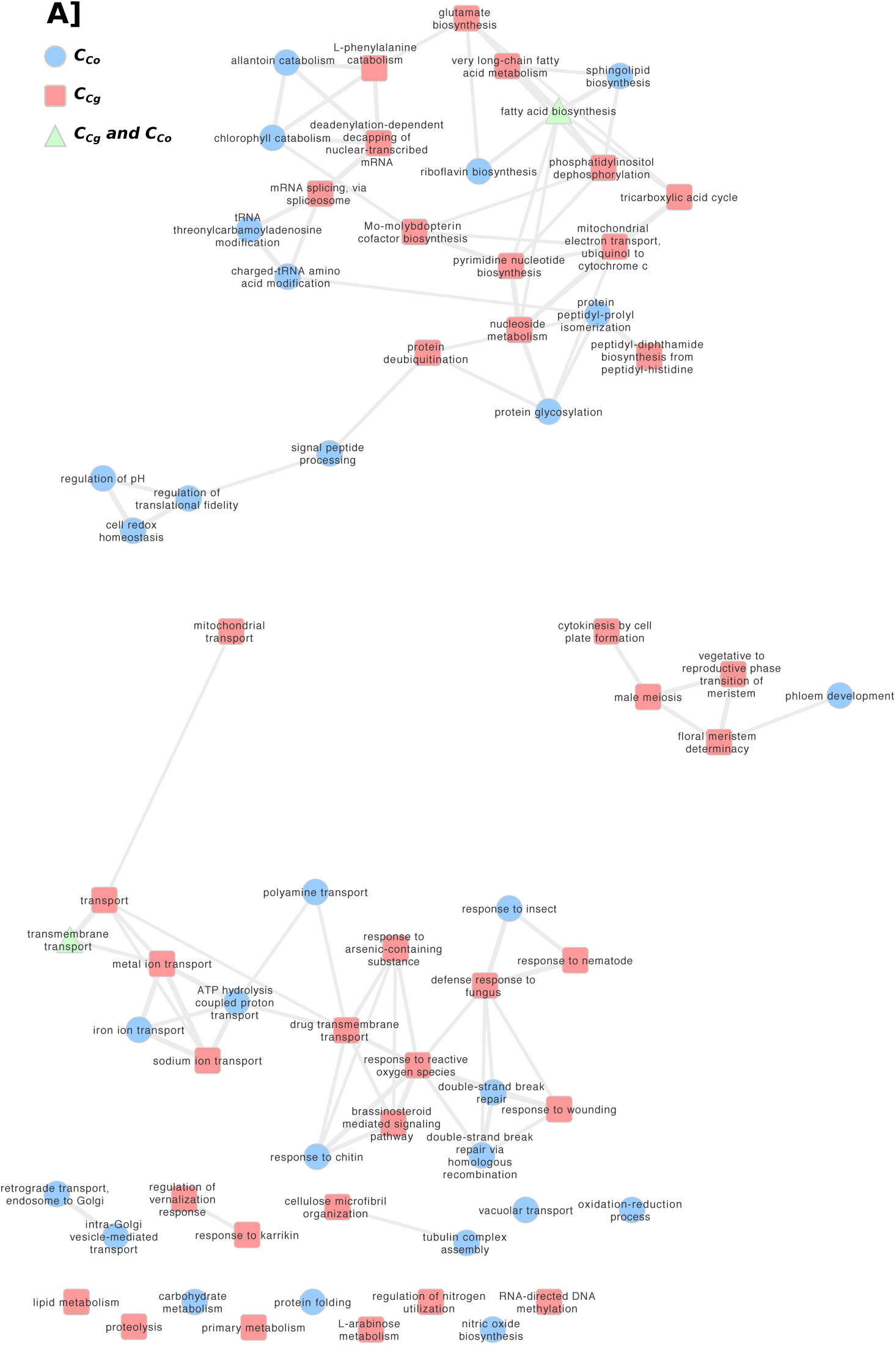
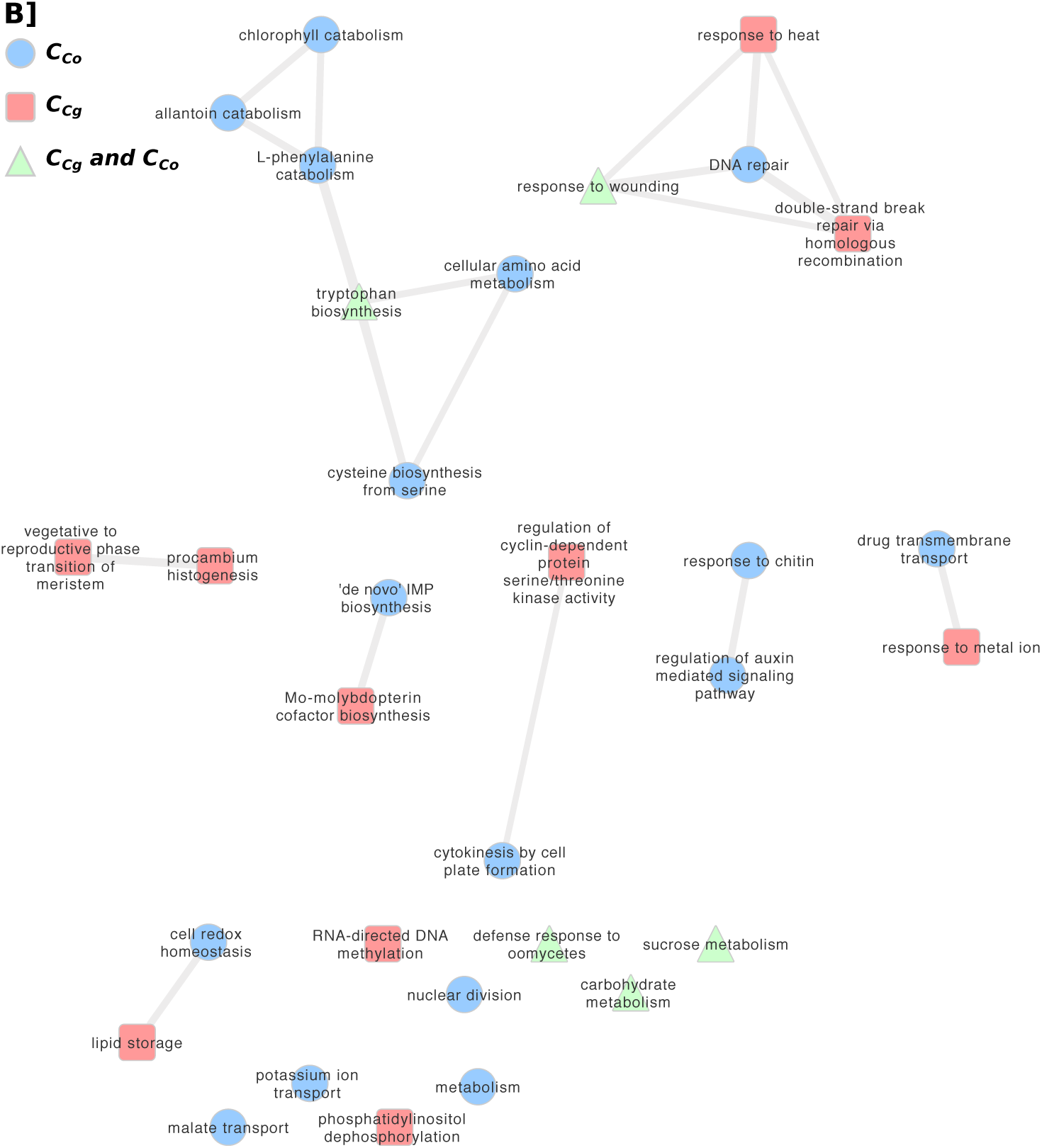
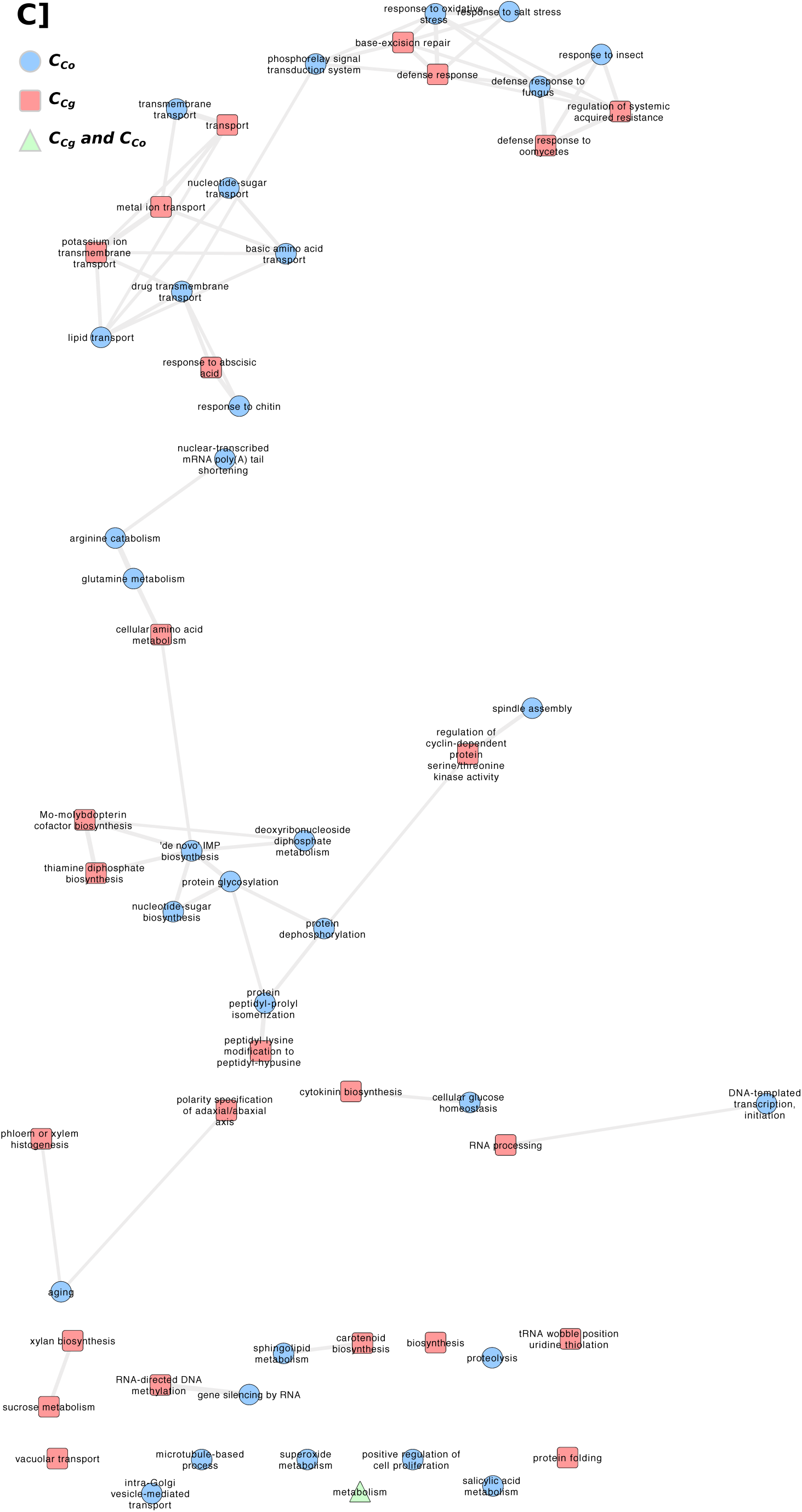
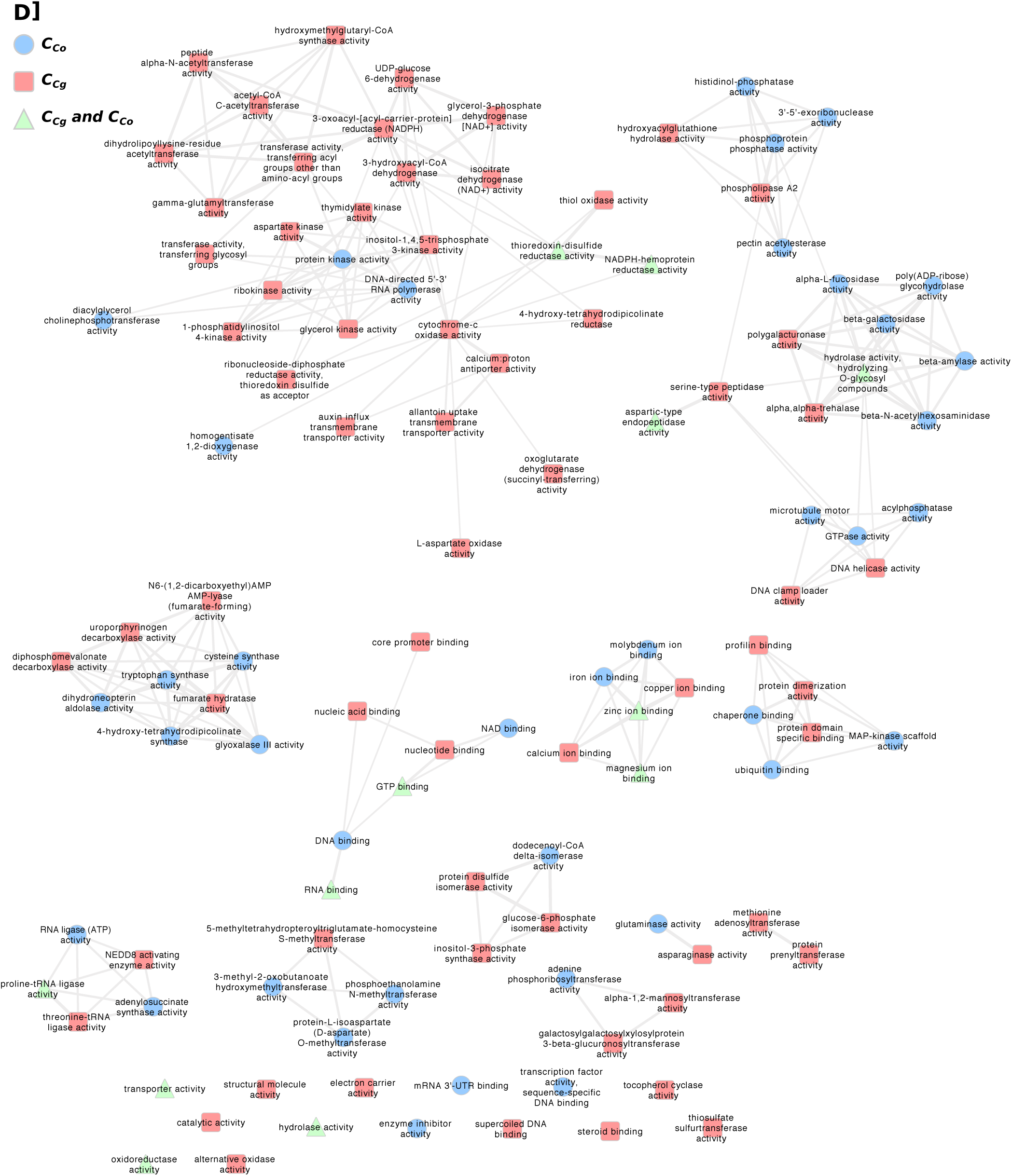
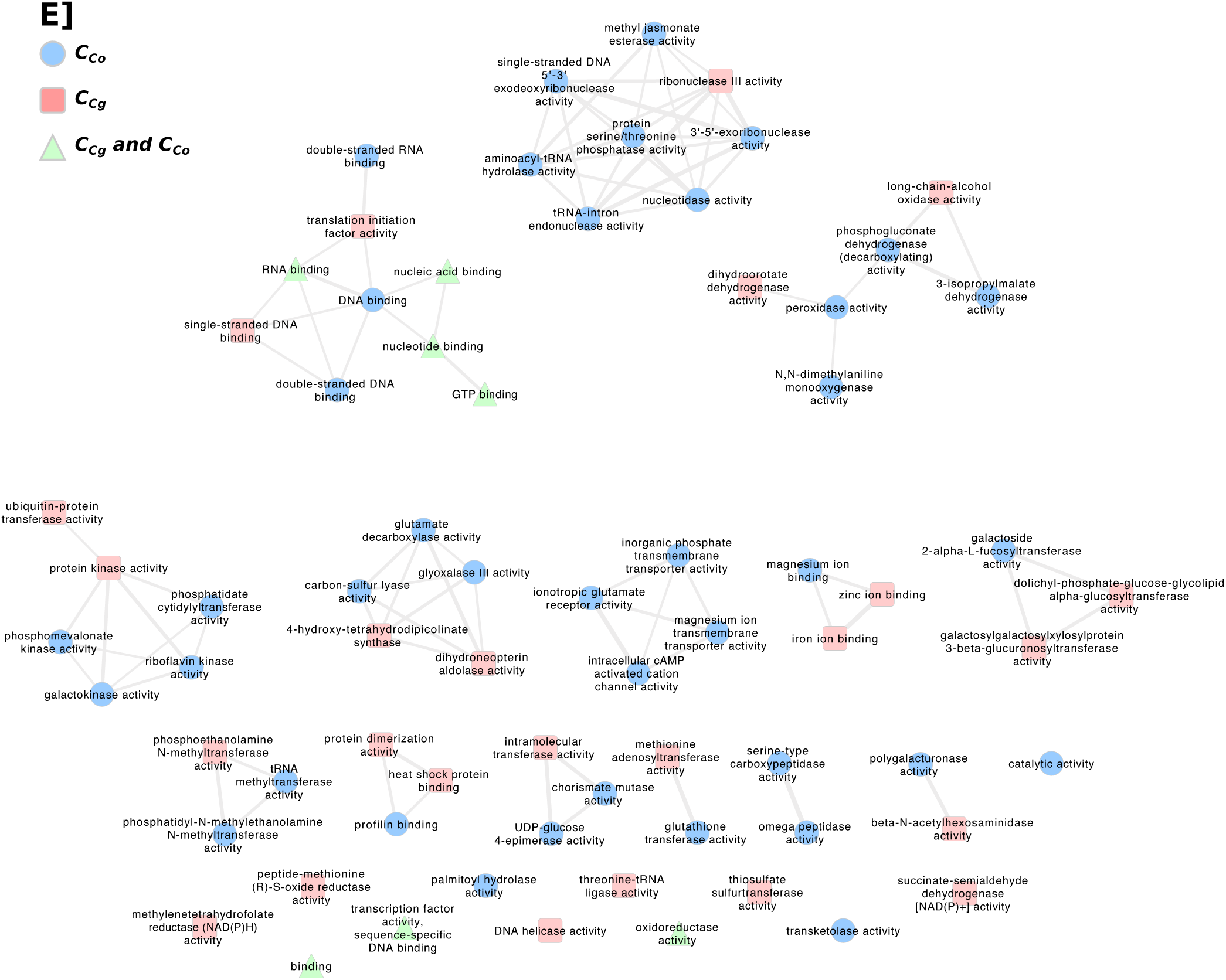
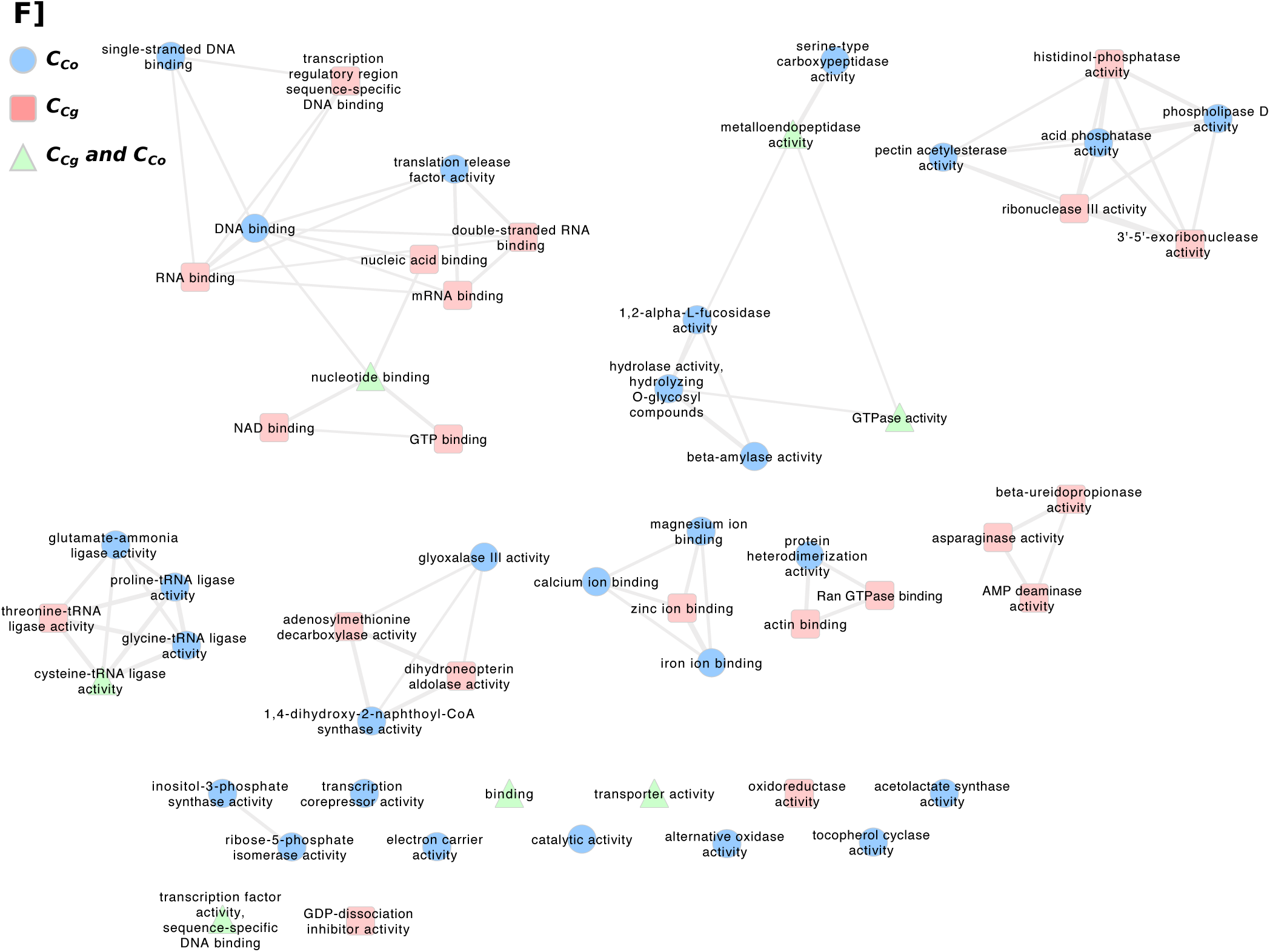
Network shared names of enriched biological processes (A, B and C) or molecular functions (D, E and F) GO term for genes showing convregence in expression between subgenomes in flowers (A and D), leaves (B and E) or roots tissues (C and F). *Co* indicates a convergence of *Cbp_Cg_*toward *Cbp_Co_*, and *Cg* corresponds to a convergence of *Cbp_Co_*toward *Cbp_Cg_*.

**Fig. S9.**
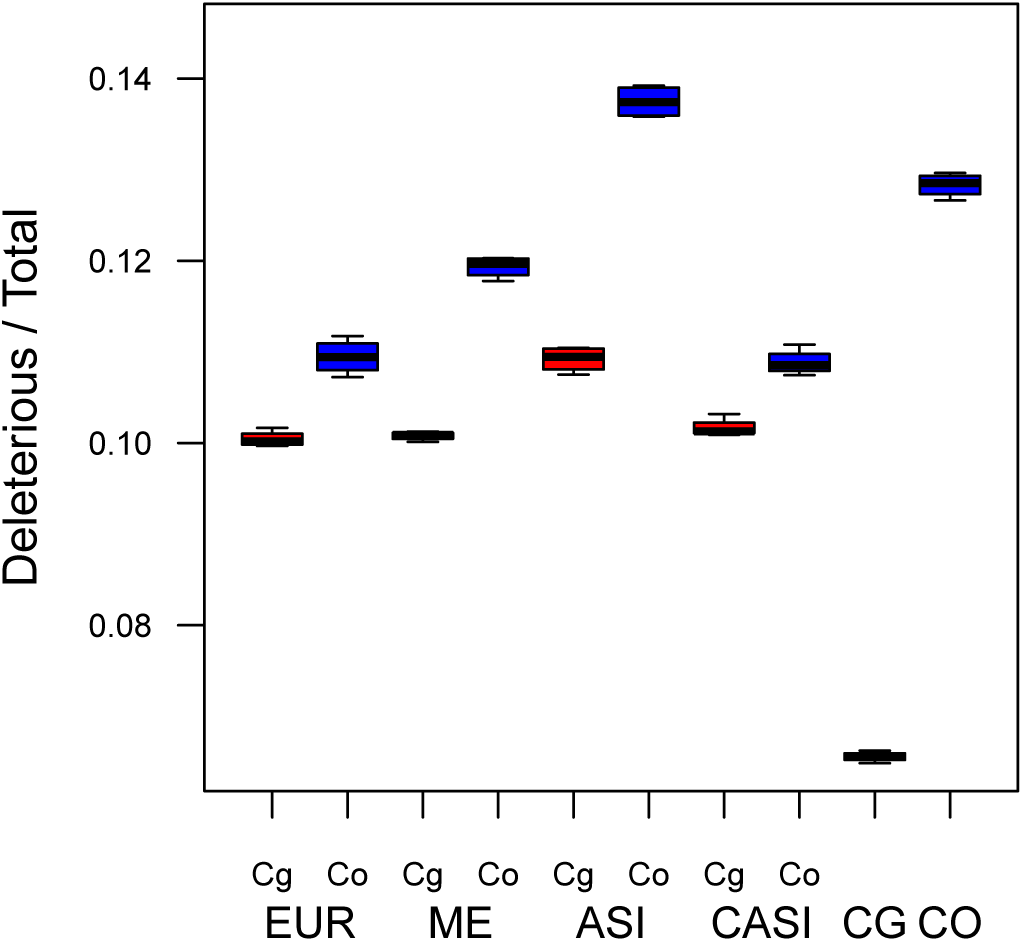
Proportion of deleterious mutations in the two subgenomes of *C. bursa-pastoris* and the genomes of its parental species. CO, CG, ASI, EUR, ME, CASI correspond to *C. orientalis*, *C. grandiflora*, and four populations of *C. bursa-pastoris*, respectively. The two subgenomes are indicated with Co and Cg. Functional effects were annotated with the *A. thaliana* SIFT database.

**Fig. S10.**
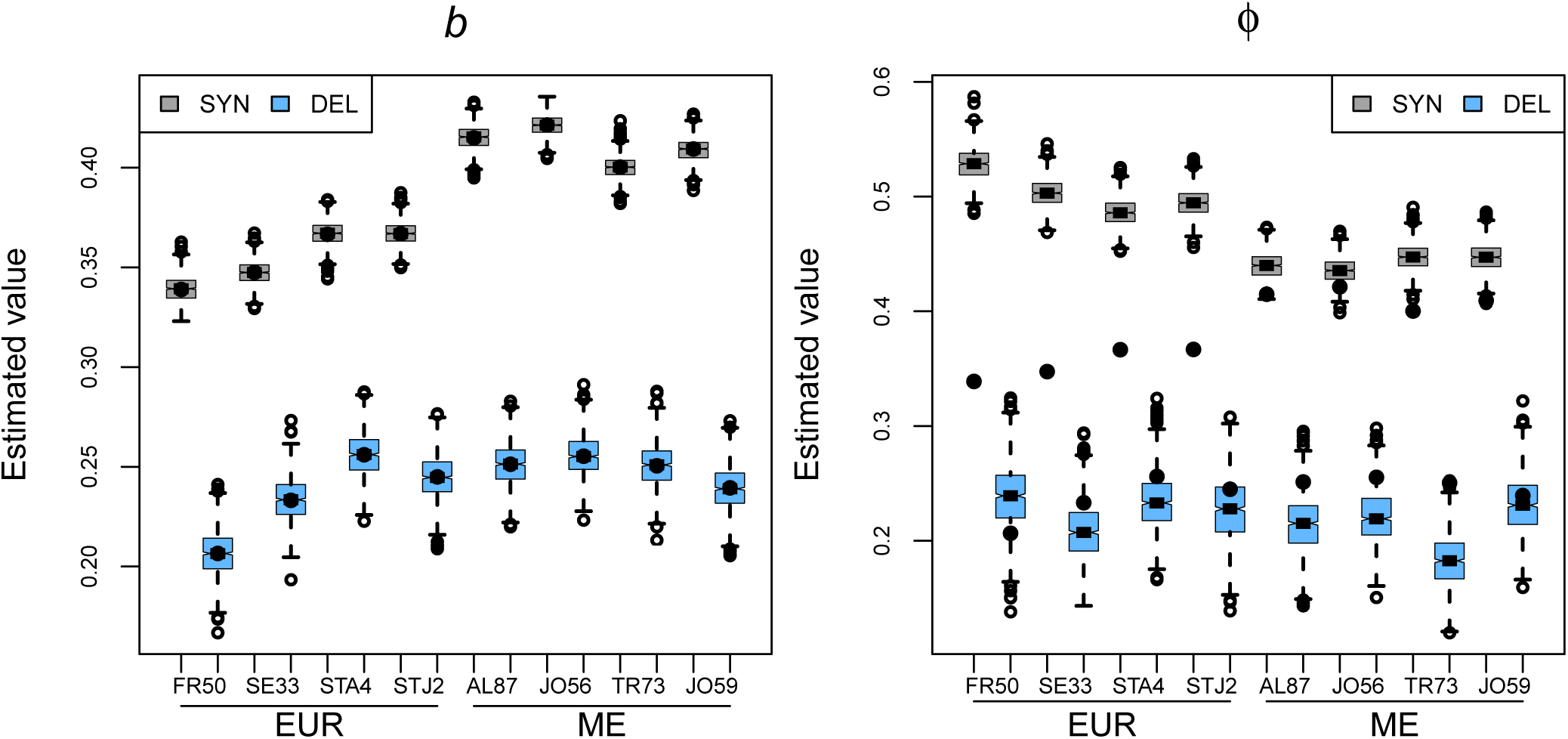
Maximum likelihood estimated parameters of the distribution of deleterious mutations on *Cbp_Cg_*genes. Each box represents the estimates of one accession, with 1000 bootstrap replicates. The estimated parameters are for synonymous mutations (SYN), and deleterious mutations (DEL). The notch of the plot represents the median and the 95% confidence interval. The black points are the point estimations with the original samples, instead of the bootstrap re-samples. The left figure shows estimates of the bias parameter, *b*, and the right figure shows estimates of the variation parameter *φ*.

**Fig. S11.**
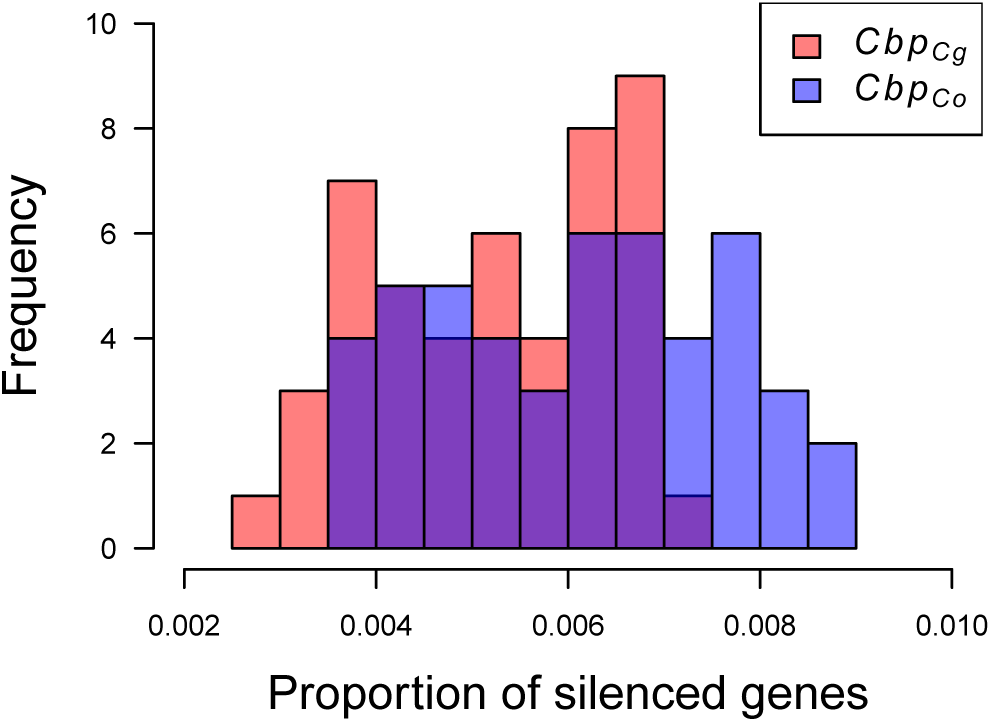
The difference in the number of silenced genes between subgenomes of *C. bursa-pastoris*.

